# Uncovering Network Architecture Using an Exact Statistical Input-Output Relation of a Neuron Model

**DOI:** 10.1101/479956

**Authors:** Safura Rashid Shomali, Seyyed Nader Rasuli, Majid Nili Ahmadabadi, Hideaki Shimazaki

## Abstract

Using observed neuronal activity, we try to unveil hidden microcircuits. A key requirement is the knowledge of statistical input-output relation of single neurons in vivo. We use a recent exact solution of spike-timing for leaky integrate-and-fire neurons under noisy inputs balanced near threshold, and construct a framework that links synaptic type/strength, and spiking nonlinearity, with statistics of neuronal activity. The framework explains structured higher-order interactions of neurons receiving common inputs under different architectures. Comparing model’s prediction with an empirical dataset of monkey V1 neurons, we find that excitatory inputs to pairs explain the observed sparse activity characterized by negative triple-wise interactions, ruling out the intuitive shared inhibition. We show that the strong interactions are in general the signature of excitatory rather than inhibitory inputs whenever spontaneous activity is low. Finally, we present a guide map that can be used to reveal the hidden motifs underlying observed interactions found in empirical data.

## Introduction

One interest in neuroscience is to *reveal in vivo neural circuitries* using recorded neuronal activities. The recent technological advances in *Connectome projects* do reveal complete wiring diagrams of certain animals (Markram et al., 2015; Oh et al., 2014; Xu et al., 2020), nonetheless, to address what computation the neural circuitry performs, it is still important to identify network architecture from *in vivo* recordings of multiple neurons. The *simultaneous intracellular recordings* are the most reliable way to identify physical connections *in vivo* (Allen et al., 2018; Arroyo et al., 2018; Gentet et al., 2010; Poulet et al., 2019; Poulet and Petersen, 2008); yet, using patch-clamp technique, one should record from neurons and all their presynaptic inputs simultaneously, to successfully find the influential synapses which form the underlying circuits, but this reliable method is limited to the small number of neurons. Instead, one could use neuronal spiking activity simultaneously recorded from a large number of neurons (Stringer et al., 2019). The *cross-correlograms* (Kobayashi et al., 2019; Perkel et al., 1967) or constructing point-process network models are *classical* approaches to infer the connectivity from spiking data (Pillow et al., 2008; Truccolo et al., 2005; Volgushev et al., 2015). However, these approaches aim at discovering connections among the recorded neurons, despite that majority of the synaptic inputs to them come from unobserved neurons. Therefore, it remains a challenge to successfully reveal the hidden neuronal circuitries, using the *activity statistics* of a limited number of neurons, *in vivo*.

The hallmark of cortical spiking activity in vivo is its *variability* (Shadlen and Newsome, 1998; Softky and Koch, 1993). It was suggested that the variability of spiking activity is the result of *balanced inputs* from excitatory and inhibitory neurons fluctuating *near spiking threshold* (Shadlen and Newsome, 1998; Vreeswijk and Sompolinsky, 1996, 1998). Such balanced inputs were confirmed by intracellular recordings of in vivo neurons (Okun and Lampl, 2008), during stimulus presentation (Tan et al., 2014). In this condition, even a moderate synaptic input can result in the spiking of the postsynaptic neuron. This, however, does not mean we can safely ignore *strong synaptic* inputs. Such *strong synaptic connections* are indeed observed in cortical and hippocampal neurons, where a *log-normal distribution* for the synaptic strength, *i*.*e*., a few strong synapses on top of a large number of weak synapses, is reported (Buzsáki and Mizuseki, 2014; Cossell et al., 2015; Lefort et al., 2009; Song et al., 2005). These influential synapses seem to act as the backbone of microcircuits. Therefore, we need to find out the architecture of these strong synapses to reveal the basic motifs of microcircuits. Previous models that link architecture to statistics of neural activity, assume *linear responses* to the synaptic input (Hu et al., 2013, 2014; Ocker et al., 2017a; Ostojic et al., 2009; Pernice et al., 2011; Rosenbaum et al., 2017; Trousdale et al., 2012) (but see Curto and Morrison 2019; Ocker et al. 2017b). However, to identify the influential inputs, the nonlinearity of input-output relation does not let us use the *linear response* methods, exclusively designed for weak synapses. There is a recent analytical solution for the Leaky Integrate and Fire (LIF) neuron, which provides the dependency of output spikes on *arbitrary synaptic input* of interest, while the effect of many weak synapses accumulates as noisy background inputs, balanced near the spiking threshold. It certifies that a strong synaptic input results in a very different and nontrivial response, compared to the weak/moderate inputs (Shomali et al., 2018).

In this study, using the aforementioned analytic solution (Shomali et al., 2018), we investigate the problem of network identification from observed correlations among spiking activity of neurons. We look at the simplest scenario: The experimentalist records spiking activity of three neurons in vivo (e.g., (Ohiorhenuan et al., 2010)), while s/he cannot directly reveal any synaptic connectivity. Will the three neurons spike independently or show correlations due to possible shared inputs? In the latter case, are such inputs shared between each pair of them, or among them all? Are shared inputs excitatory or inhibitory? And finally, does either of the three observed neurons make any direct synaptic connection to another of them?

We obtain *pairwise* and *triple-wise* interactions (Amari, 2009a; Nakahara and Amari, 2002) in the simultaneous spiking activity of three LIF neurons, and compare them with experimentally observed results of Ohiorhenuan et al. in monkey V1. They found significant positive pairwise and negative *triple-wise interactions* for spatially close neurons (Ohiorhenuan et al., 2010; Ohiorhenuan and Victor, 2011). Negative triple-wise interactions, observed in cortical and hippocampal neurons (Ohiorhenuan et al., 2010; Shimazaki et al., 2015; Yu et al., 2011), indicate a significantly higher probability of simultaneous silence among three neurons than those expected from their rates and pairwise correlations. Intuitively, excess simultaneous silence can be induced by suppression of these neurons caused by common inhibitory inputs. However, the analytical input-output relation quantitatively reveals that a non-intuitive architecture of *common excitatory inputs, each shared by a pair of neurons* (excitatory-to-pairs), does explain the observed negative triple-wise interactions, and rules out shared inhibition. We investigate the robustness of our results to sub/suprathreshold regimes, directional/recurrent connections among observed neurons, and to adaptative neurons. We confirm many of these results, particularly the significance of excitatory inputs to pairs’ motif, remain intact. Plotting triple-wise versus pairwise interactions, for all basic motifs, we analytically provide a 2D guide map; each motif occupies its own region there. This guide map can be used to identify the hidden motifs underlying observed interactions found in empirical data.

## Results

### Spike probability of a leaky integrate-and-fire neuron: Near threshold regime

First, we introduce the statistical properties of our cortical neuron model operating under in-vivo like conditions (Shomali et al., 2018). We evaluate the probability of spiking within a given time window; it becomes the building block to construct the population activity of such neurons. To this end, we begin with a Leaky Integrate and Fire (LIF) *postsynaptic neuron* with membrane’s time constant of *τ*_m_, and a resting potential of *V*_r_:

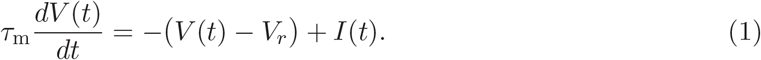

The neuron spikes when its membrane potential, *V* (*t*), hits the spiking threshold, *V*_*θ*_; then *V* (*t*) resets to *V*_*r*_. The input current of *I*(*t*), also, consists of two parts: (a) a transient signaling input which represents the input from the influential synapses with arbitrary strength, Δ*I*(*t, A, τ*_*b*_), and (b) the effect of all other independent presynaptic inputs accumulated as a *fluctuating background input, I*_0_(*t*):

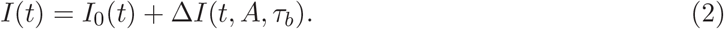

We model the fluctuating background input as Gaussian white noise, so it well replicates the synaptic inputs to V1 neurons when a visual stimulus is presented (Tan et al., 2014). Conclusively, *I*_0_(*t*) has a mean drive of *Ī* and a variance of 2*D/τ*_m_; here the diffusion coefficient of *D* measures *I*_0_(*t*)’s level of noise. The signaling input, Δ*I*(*t, A, τ*_*b*_), is characterized by its amplitude (or efficacy), *A*; and its arrival time, *τ*_*b*_.

The fluctuating *I*_0_(*t*), is one important source of *variability*; its *stochastic* nature, however, makes it impossible to solve Eq.(1) and find the exact spike-time *deterministically*. Thus, people try to address the *probability of spiking* (Brunel and Hakim, 1999; Gerstner et al., 2014a; Shomali et al., 2018). Their essential mathematical tool is the *Fokker-Planck* (or diffusion) equations (Risken and Eberly, 1985), which addresses the probability density that postsynaptic neuron spikes at time *t*, given it had a known value of membrane potential at the initial time of *t*_0_. However, even in the absence of any signaling input, the corresponding Fokker-Planck equation has not been solved yet. There exists an analytical solution but for a very specific case *Ī* = *V*_*θ*_ (Bulsara et al., 1996; Wang and Uhlenbeck, 1945), which is known as the threshold regime representing a physiologically plausible situation for in vivo neurons. Recently, Shomali et al. were able to extend that analytic solution for spike density when signaling inputs arrive on top of background noise (Shomali et al., 2018). They considered a near threshold neuron, *Ī* ≃ *V*_*θ*_, that receives a transient signaling input (*i*.*e*., the synaptic time constant of *τ*_s_ is smaller enough than the membrane time constant, *τ*_m_). They solved the Fokker-Planck equation, and *analytically* found the probability density of spiking (known also as Inter-Spike Interval distribution, ISI) for arbitrary strength and shape of the signaling input.

Using that framework, we assess the effect of the signaling input on the activity of postsynaptic neurons (Fig. 1A, right). We ask two successive questions: First, what is the probability density of a spike occurrence at time *τ* after signal arrival, *f* (*τ*)? And second, what is the probability of observing one or more spikes in a time window of Δ, after the occurrence of presynaptic signal with a strength *A, F*_*A*_(Δ)? Figure 1 depicts our results for these two questions (see Methods for the detailed derivation of *f* (*τ*) and *F*_*A*_(Δ)). Particularly, *F*_*A*_(Δ) is the predictable quantity that relates the neuron model to the observed spiking activity of neurons.

**Fig 1.**
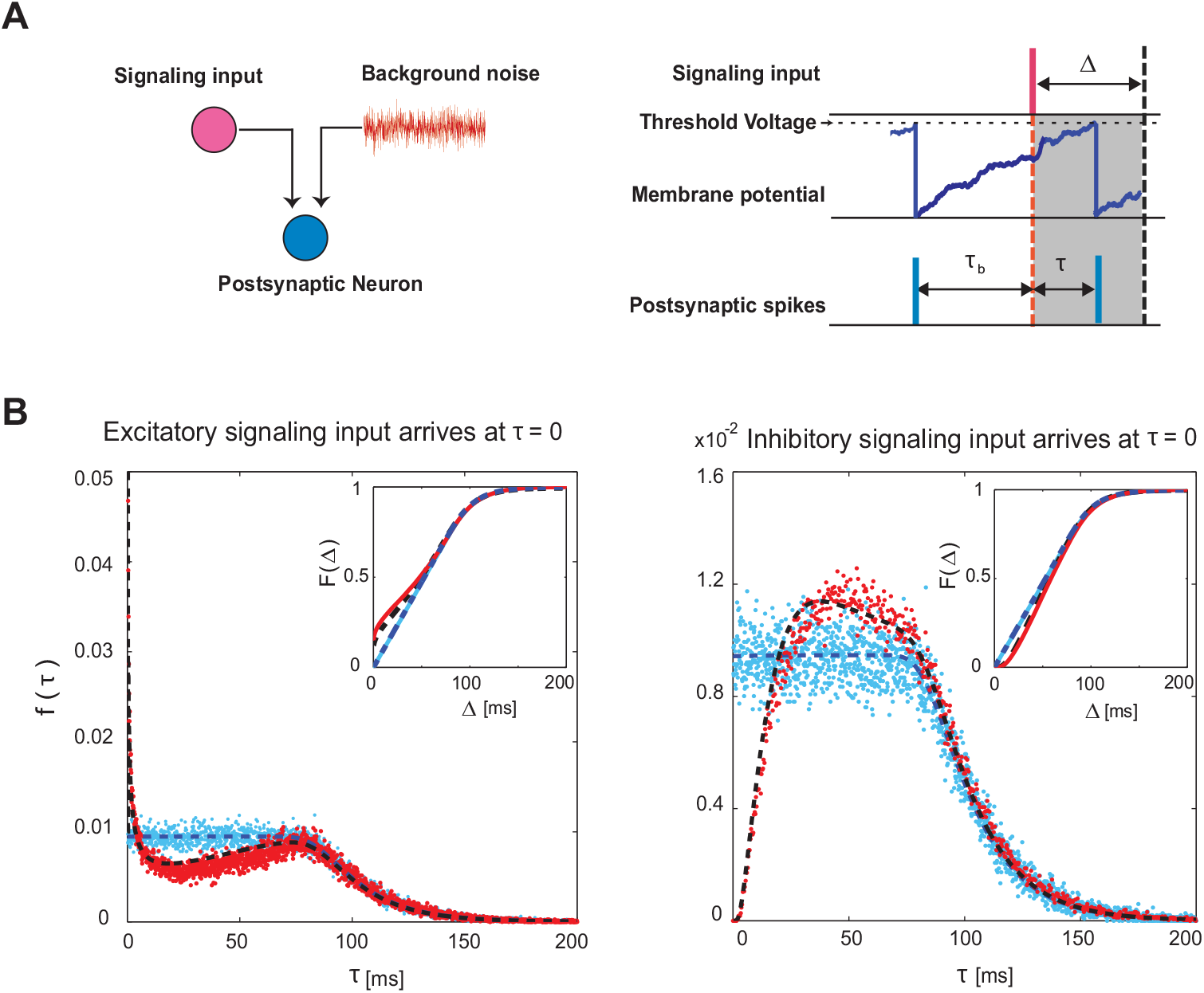
Analysis of a postsynaptic neuron receiving signaling input on top of background noise. **(A)** Left: A schematic model of a postsynaptic neuron driven by background Gaussian noise and a transient signaling input. Right: Timing of postsynaptic spikes and the arrival time of the signaling input. The postsynaptic neuron generates a spike (blue tick) then its membrane potential resets. While the potential rises, a signaling input (red tick) arrives at *τ*_*b*_ after the last spike, which changes the trajectory of the membrane potential. The postsynaptic spike occurs at *τ* after arrival time of the signaling input. The gray shaded area of width Δ indicates the observation time window during which the probability of postsynaptic activity pattern is computed. **(B)** The probability density of the first spike at time *τ* after arrival (Eq. 11) of excitatory (left) and inhibitory (right) signaling input. The simulation results (red dots) and the analytical solutions (dashed black line, Eq. 11) are well matched. The same results for a signaling input with zero amplitude are depicted. Inset: The probability of spiking within Δ after signaling input arrival, Eq. 12. The parameters are: *V*_*θ*_ = 20 mV, *τ*_m_ = 20 ms, *A* = 5 mVms, and the diffusion coefficient is *D* = 0.74 (mV)^2^ms. The values of the parameters are chosen from the physiologically plausible range (McCormick et al., 1985).

Figure 1B shows the marginalized spiking density at time *τ* after signaling input arrival for the square shape input (Eq. 9, dashed black lines). Early spiking, i.e., small *τ*, after the arrival of excitatory (or inhibitory) input is much more (or less) probable, compared with no-signaling input case (dashed blue lines). However, the spiking densities with and without the signaling input are virtually identical at sufficiently large *τ*; indicating the short-lasting effect of the signaling input. Accordingly, the cumulative distribution functions, *F*_*A*_(Δ), (Fig. 1B insets) with and without the signaling input differ for small Δ, but are indistinguishable for large Δ. This result implies that we cannot discern the presence of signaling input if we use a large time window.

Using *F*_*A*_(Δ), one can find the probability of various spiking patterns for multiple neurons receiving common signaling inputs, under the assumption that the neurons are conditionally independent. This calculation is described in the next section.

### Spike density of in-vivo LIF neurons can be used to model population activity driven by common inputs

We provide a framework for computing the statistical properties of the population activity of the LIF neurons that receive common excitatory or inhibitory inputs (Fig. 2). We investigate both pairwise and triple-wise interactions among neurons while they are receiving common inputs on top of noise in basic possible motifs. Firstly, we consider only two neurons for simplicity, whereas the framework will be extended to three neurons in the next section. Suppose that the two postsynaptic neurons receive a common signaling input in addition to independent background noise. Figure 2 illustrates the timing of the postsynaptic spikes before and after the common input arrival. By binning their spike sequences with a Δ time window (Fig. 2, middle, and left), we compute the probabilities of activity patterns of postsynaptic neurons. We assume that neurons sparsely receive a random common input with firing rate *λ* in a Poissonian fashion. We then, segment the spike sequences using bins aligned at the onset of common input. Let *x*_*i*_ = {0, 1} (*i* = 1, 2) be a binary variable, where *x*_*i*_ = 1 means that the *i*th neuron emitted one or more spikes in the bin, while *x*_*i*_ = 0 means that the neuron is silent.

**Fig 2.**
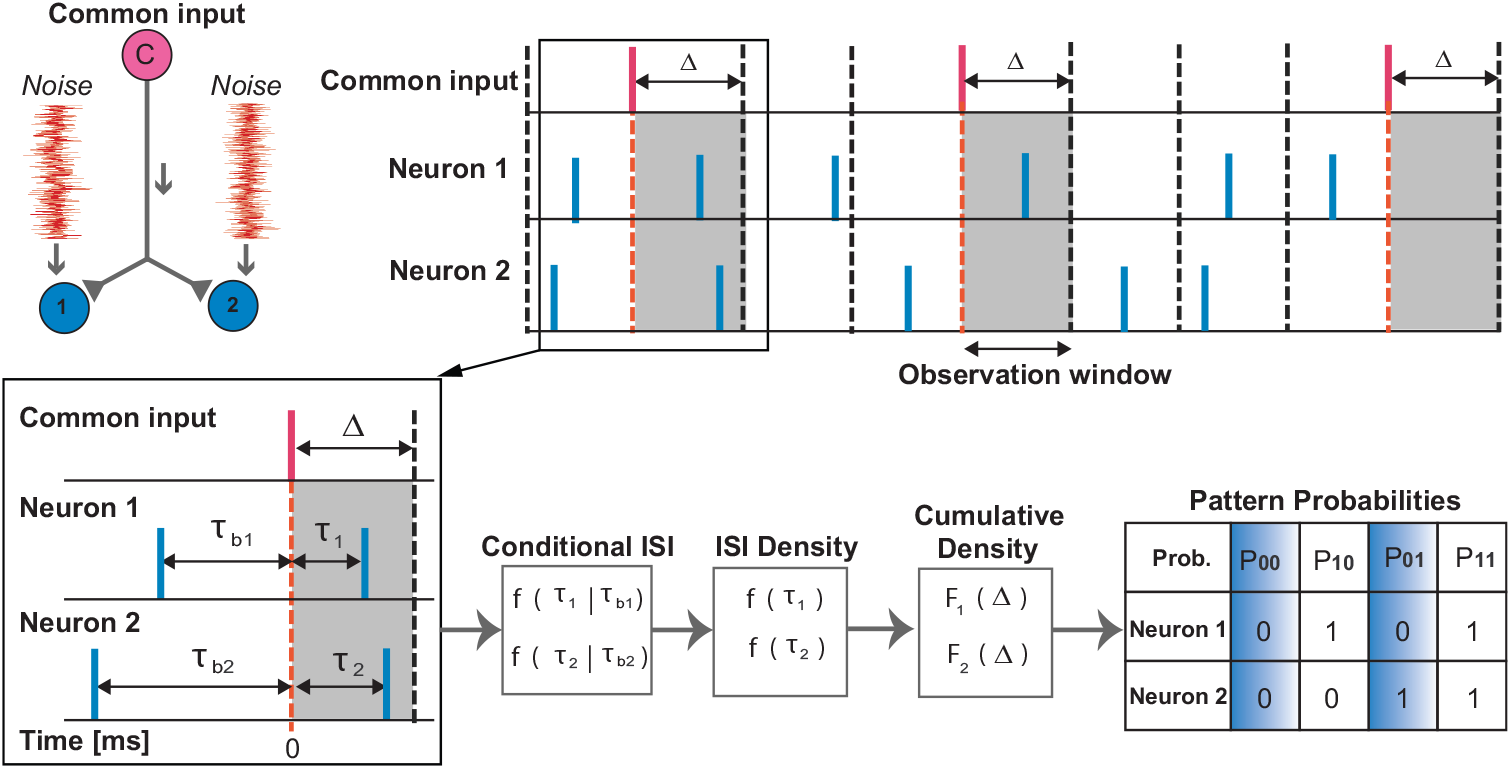
Analysis of two neurons receiving common signaling input on top of background noise. Top, left: A schematic model of two postsynaptic neurons (Neuron 1 and Neuron 2, blue circles) driven by independent noise and a common signaling input (pink circle). Top, middle and right: Spike trains of two postsynaptic neurons (blue spikes) receiving common signaling inputs (red spikes) with rate *λ*. The gray shaded area of width Δ indicates the time window during which we compute the probabilities of postsynaptic activity patterns. Bottom, left: Timing of postsynaptic spikes relative to the arrival time of the common input. The last spike of Neuron 1 (Neuron 2) has occurred *τ*_*b*1_ (*τ*_*b*2_) before the arrival of common input, and their next spikes happen at *τ*_1_ (*τ*_2_). Bottom, middle: The conditional ISI density after input arrival is calculated by Eq. 10. Marginalizing over previous spike (*τ*_*b*_), one obtains the probability of spiking after input arrival (ISI density, Eq. 11). The next step is to calculate the cumulative distribution function (Eq. 12) that is the probability of having one or more spikes within window Δ. Bottom, right: Based on the cumulative distribution function and the fact that neurons are conditionally independent, the probability of having a particular pattern of spikes for two neurons is obtained. Four possible binary activity patterns (00, 01, 10, 11) of two postsynaptic neurons and their associated probabilities (*P*_*ij*_, *i, j* ∈ {0, 1}). The ‘1’ denotes the occurrence of at least one spike within the Δ time window, whereas the ‘0’ represents the silence of the neuron within this window.

To formally investigate the neuronal correlation, we use the information-geometric measure of interaction, *θ*_12_ (Amari, 2009b; Martignon et al., 2000; Nakahara and Amari, 2002; Tatsuno and Okada, 2004) (Eq. 14 in Methods). This information-geometric measure of the correlation (SVI) is recommended over the classical covariance or correlation coefficient (Amari, 2009a; Martignon et al., 2000; Nakahara and Amari, 2002; Tatsuno and Okada, 2004) because this measure is not affected by the estimated firing rates of neurons while other measures are influenced by firing rates (i.e, this method is invariant or orthogonal to firing rates in terms of the Fisher metric, see SVI). For two neurons, the probabilities of the activity patterns can be constructed using the probability of spike occurrence in a bin Δ (Eq. 12, Fig. 2). Since two neurons receive common inputs, they are conditionally independent. Therefore, the probabilities of activity patterns in a bin are given by:

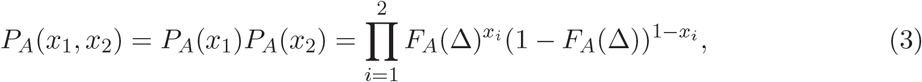

where *A* stands for the amplitude of the common input current (*A* = 0 represents the absence of common input). The rate of common input (*λ*) is applied at *λ*Δ × 100% of the bins whereas it is absent in (1 −*λ*Δ) × 100% of the bins. Accordingly, we consider the spike sequences of two postsynaptic neurons as a mixture of the two conditions: Neurons either receive (*A* ≠ 0) or do not receive (*A* = 0) common input. Hence the probabilities of the activity patterns are given by

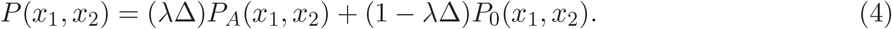

From this probability mass function, Eq. 4, one can compute the neuron’s pairwise interaction, denoted as *θ*_12_ (Eq. 14). This mixture model provides approximated probabilities for the activity patterns of the LIF neurons. This is an approximation of the actual dynamics because, if a neuron does not spike within Δ [ms] after the common input, the effect of the augmented membrane potential is carried over to the next bin, and then the binary activities are no longer a simple mixture of the two conditions. Such situations should happen often if the bin size is small compared to the mean postsynaptic inter-spike interval. So we test if this approximation predicts the interaction in the parallel sequences of the two LIF neurons, and examine reasonable bin sizes. We compare the pairwise interaction predicted by the mixture model with the simulated spike sequences in Figure 3A. It displays the interaction for different bin sizes, Δ, when the two neurons receive common excitatory input. The pairwise interaction predicted by the mixture model agrees with the simulation results (left panel, red and gray lines, respectively). The result also shows *θ*_12_ increases with the rate of common input (Fig. 3A, Right). However, if we increase the bin size, the probability of having one or more spikes within Δ increases and saturates to 1 (Fig. 1B, Inset) regardless of the presence or absence of the signaling input. This means *F*_*A*_(Δ)*/F*_0_(Δ) → 1, which results in vanishing pairwise interaction as the bin size increases. Therefore, we can hardly discriminate between the presence and absence of common input if we use a large bin size.

**Fig 3.**
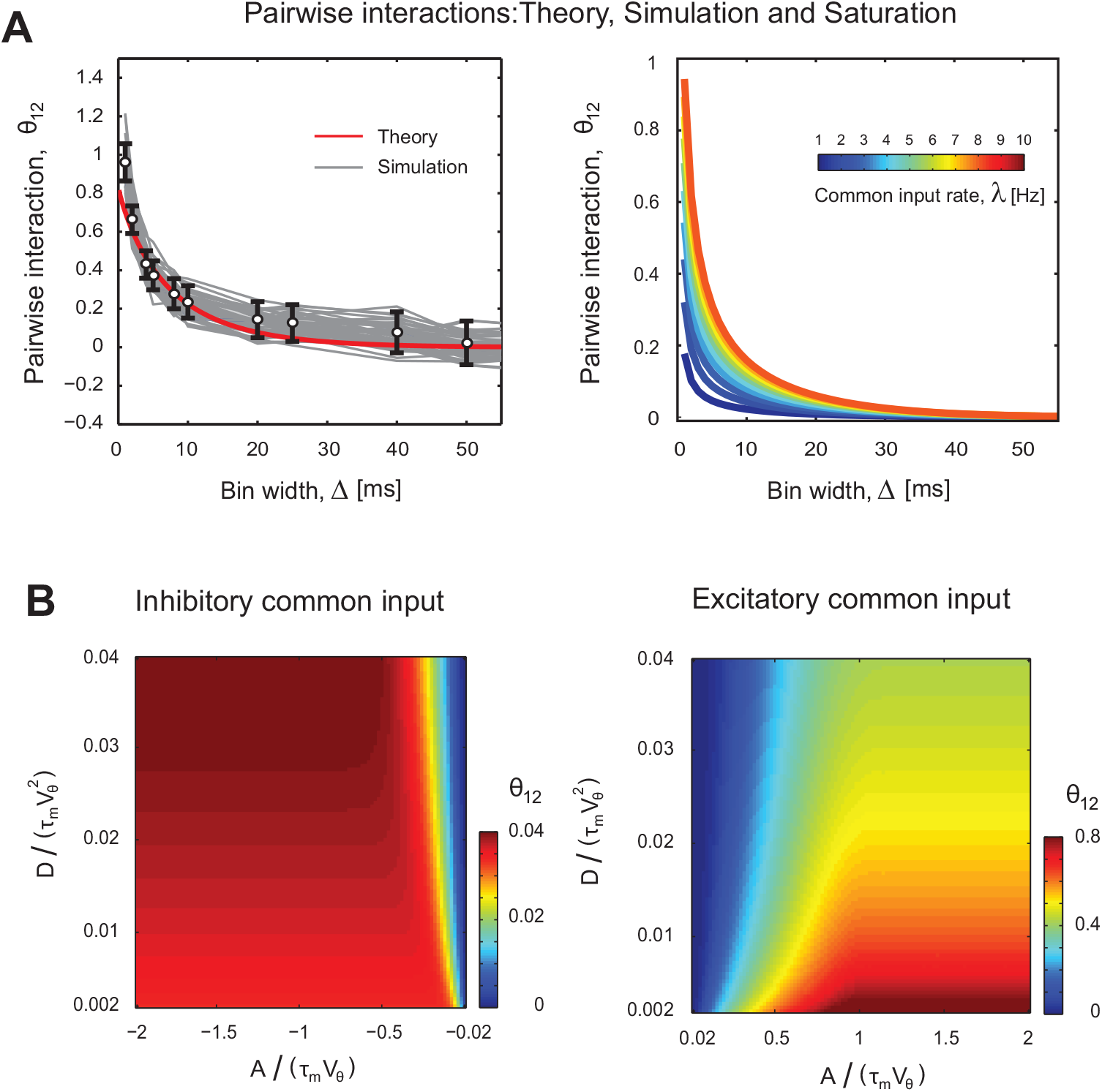
Analysis of pairwise interaction of two neurons receiving common signaling input on top of background noise: **(A)** The interaction (*θ*_12_, Eq. 14) of two postsynaptic neurons as a function of bin size, Δ. We use a physiologically plausible range of parameters as: *V*_*θ*_ = 20 mV, *τ*_m_ = 20 ms, *A* = 5 mVms, and the diffusion coefficient *D* = 0.74 (mV)^2^ms (McCormick et al., 1985). Left: The pairwise interaction computed from simulated spike sequences (gray lines: 50 individual trials each containing about 2500 spike occurrence of common input; dots and error bars: mean± standard deviation) compared with the analytic result of the mixture model (red line, Eq. 14) for common input rate *λ* = 5 Hz. Right: Analytical value of *θ*_12_ as a function of Δ for different common input rates, *λ*. **(B)** The pairwise interaction as a function of the scaled diffusion coefficient, 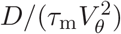, and the shared signal strength, *A/*(*τ*_m_*V*_*θ*_), for excitatory (right) and inhibitory (left) common inputs, with *λ* = 5 Hz and Δ = 5 ms.

We examine the pairwise interactions by changing two independent parameters, the scaled amplitude of the signaling input *A/*(*τ*_m_*V*_*θ*_) and the scaled variability of the noisy background input 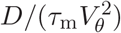 (Fig. 3B). As expected, the pairwise interactions are positive for both common excitatory and inhibitory inputs. However, the interactions are significantly weaker in the inhibitory case. This indicates that it is difficult to observe the effect of common inhibitory input for this range of postsynaptic firing rates, and that the strong pairwise interactions are the indicator of having common excitatory inputs.

For each value of normalized diffusion coefficient (level of inputs’ noise), 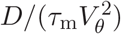, there exists a critical normalized amplitude for common *excitatory* input, *A/*(*τ*_m_*V*_*θ*_) ∼ 1. Upward this critical value, the postsynaptic neuron’s spiking density, and consequently pairwise interaction, does not change anymore (Fig. 3B, right). The saturation value of pairwise interaction is inversely correlated with the normalized diffusion coefficient; since higher normalized diffusion coefficient (level of the noise) disperses the voltage of the membrane, the probability of spiking after common input arrival decreases. Similar behavior for *θ*_12_ is observed for inhibitory input (Fig. 3B, left). Nevertheless, in contrast to the common excitatory input case, pairwise interaction and 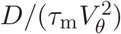 are directly correlated.

### Higher-order interaction of three neurons depends on types of common inputs and network architecture

We now extend the analysis of neural interactions to three neurons. The motivation to investigate the interactions among three neurons comes from experimental studies (Ohiorhenuan et al., 2010; Ohiorhenuan and Victor, 2011) that investigated the activities of three neurons simultaneously (Fig. 4A). For two neurons, there is one possible shared input’s architecture: A common input to both of them. For three neurons to induce correlation among them, however, it can be either *(i)* a shared input among three of them (red connections in Fig. 4A), or (*ii*) one or more shared inputs to each pair among them (green connections). Assuming symmetry, the former one leads to a *star architecture* or a “common input to trio” (Fig. 4A, middle) while the latter one makes a *triangle architecture* or “common inputs to pairs” (Fig. 4A, right).

**Fig 4.**
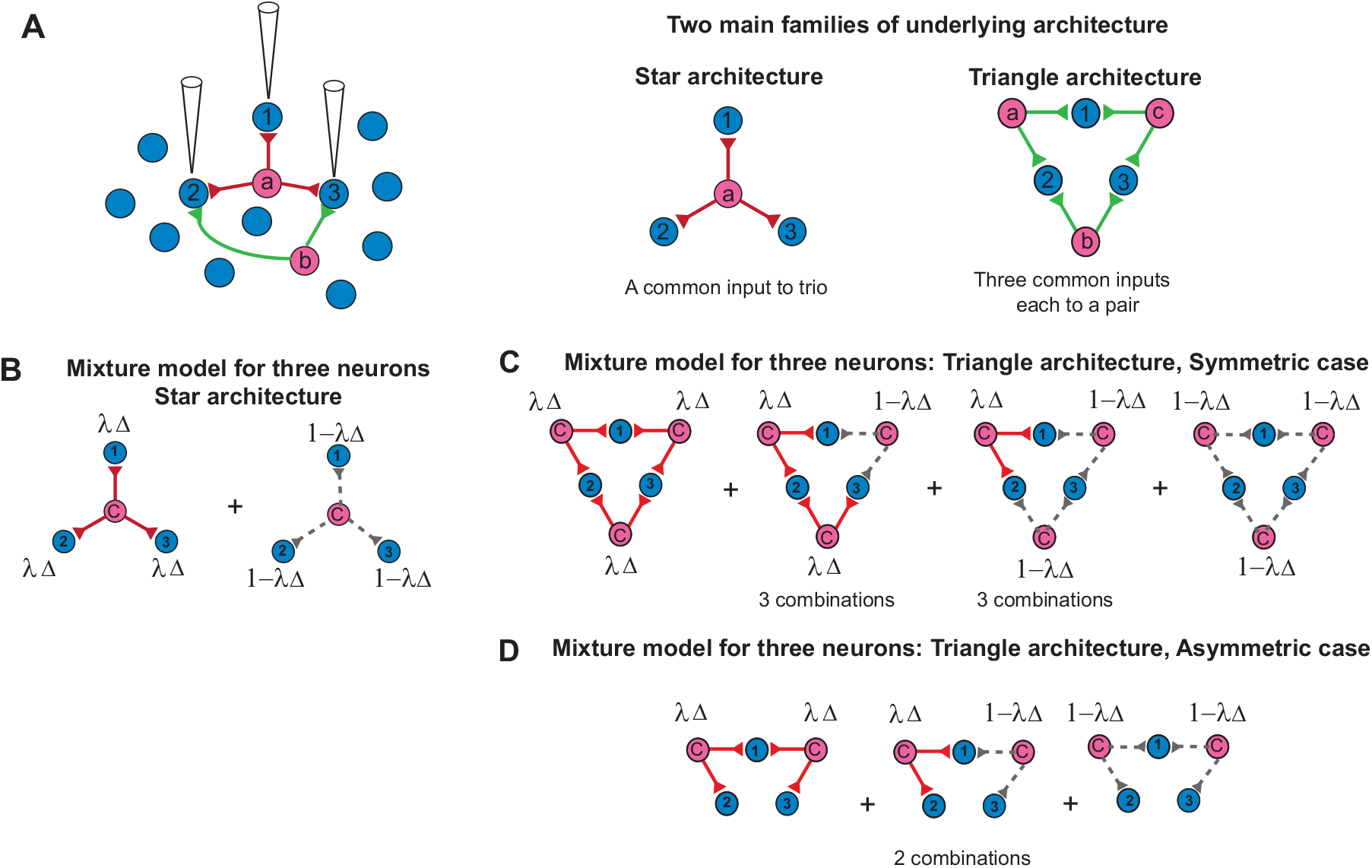
**(A)** A schematic of simultaneous recording of three postsynaptic neurons. The neurons (blue circles) operate independently in the absence of any *common input*. A common input (pink circle) can be an input to three of them (red connections), or at least two of them (green connections). Right: Assuming symmetric architectures, we have two main families: One common input to trio (star architecture, left), or three common inputs each to a pair (triangle architecture, right). **(B)** A mixture model for three neurons in the star architecture is composed of two conditions in which the neurons receive the common input (red lines) with probability *λ*Δ and they do not receive it (gray dashed lines) with probability 1 −*λ*Δ. **(C)** A mixture model for three neurons with the symmetric triangle architecture is composed of four distinct conditions that arise by the combination of the presence or absence of each common input (see main text). **(D)** For asymmetric triangle architecture, the number of possible cases is reduced to three.

For investigation the neuronal correlation among three neurons, we use the information-geometric measure of a triple-wise interaction, *θ*_123_ (Amari, 2009b; Martignon et al., 2000; Nakahara and Amari, 2002) (Eq. 16 in Methods). As the pairwise interaction extracted a pure interaction of two neurons, this triple-wise interaction measure is also not affected by firing rates and joint firing rates of two neurons, and extracts a pure triple-wise effect that can not be inferred from the first and second-order statistics of the population (SVI). There are two basic motifs that can induce triple-wise interaction among neurons (Fig. 4A, middle and right) as described below.

#### I. Common input is given to three neurons: Star architecture

In star architecture, three neurons receive a single common signaling input simultaneously (Fig. 4B). The conditional probability of activity patterns when a common input generates a spike given to all three neurons with probability *λ*Δ (red lines) is 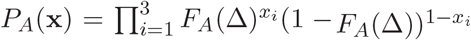, where **x** = (*x*_1_, *x*_2_, *x*_3_) is the spiking activity for three neurons. Similarly, the probability mass function for three neurons receiving no common input with probability 1 −*λ*Δ (gray dashed lines), is obtained by 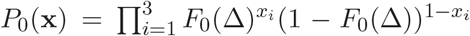 Thus, we model spike occurrence as a mixture of the two conditions in which neurons receive and do not receive common input:

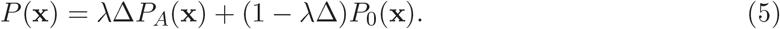

From this probability mass function, we can compute the triple-wise interaction of three neurons according to Eq. 16.

#### II. Common inputs are given to pairs of three neurons: Triangle architecture

In triangular architecture, we assume that each pair of three neurons receives common signaling input from an independent presynaptic neuron with frequency *λ* (Fig. 4C). The first, second, and third common inputs project to neurons 1 and 2, neurons 2 and 3, and neurons 1 and 3, respectively (symmetric case). The three common inputs are independent, and occur with equal frequency, *λ*. By taking into account the occurrence probabilities, one could obtain the mixture model. The resulting mixture models are given in Methods, which also include the asymmetric common input architecture in which there are only two common inputs out of three (asymmetric case) (Fig. 4D).

The triple-wise interaction parameters computed from the simulated spike sequences of post-synaptic neurons are compared with the theoretical predictions, using the mixture models (Fig. 5A and B, left). The activities of neurons that receive a simultaneous common excitatory input (star architecture) are characterized by positive triple-wise interactions (Fig. 5A, left) whereas the activities of neurons that receive independent common excitatory inputs to pairs (triangular architecture) are characterized by negative triple-wise interactions (Fig. 5B, left). Figure 5A and B, right, show that triple-wise interaction decreases as the bin size increases for the same reason given for pairwise interaction (Fig. 3A, right). The dependency of triple-wise interaction on the common input rate is also shown in the right panels in Figure 5A and Figure 5B.

**Fig 5.**
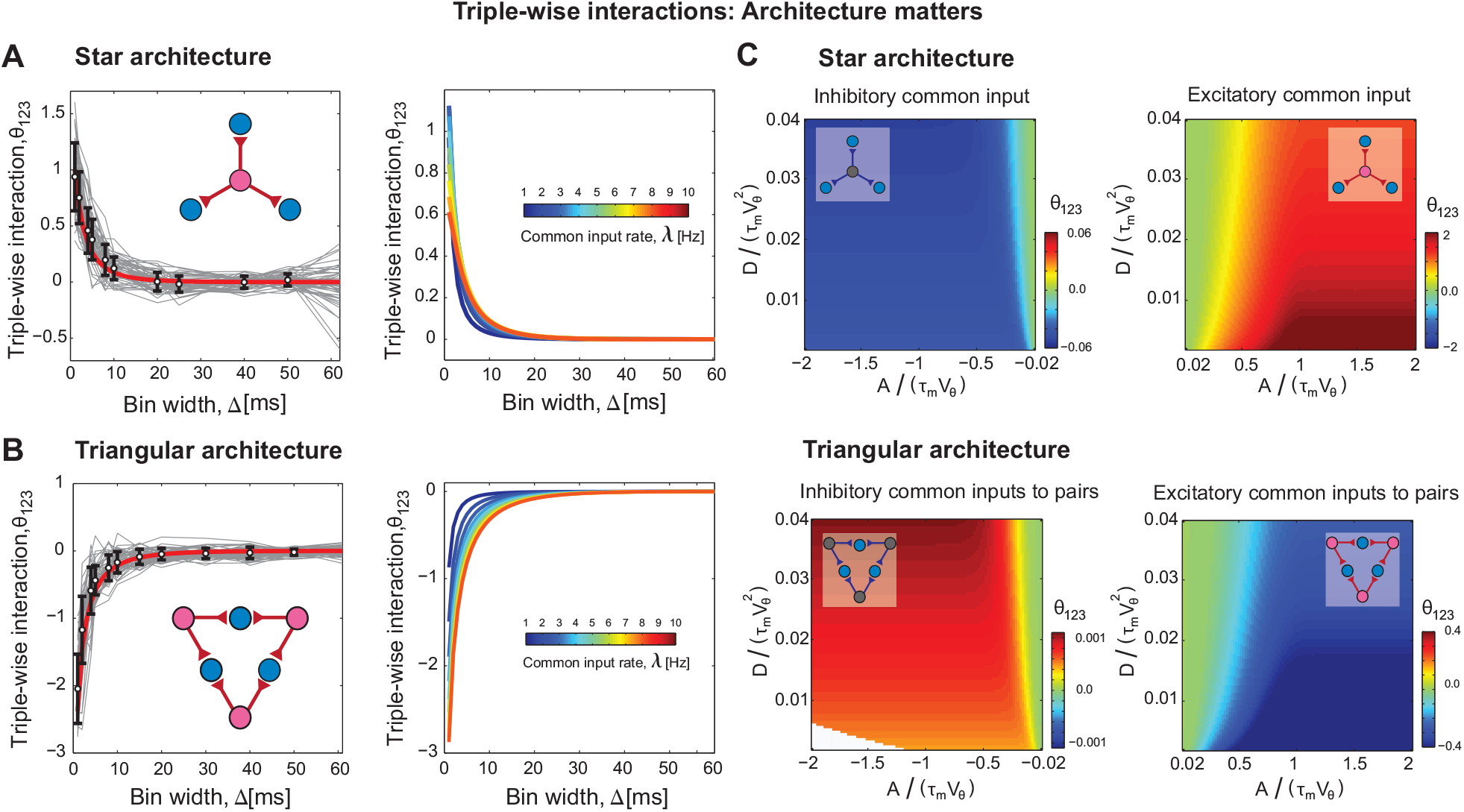
Comparison of the *triple-wise interaction* of 3 **LIF** neurons, *θ*_123_, in the two leading architectures: **(A)** The interaction when three postsynaptic neurons, blue ones, are in a *star architecture* receiving a common excitatory signal *A* = 5 mV ms, simultaneously. The parameters are: *τ* = 20 ms, *V*_*θ*_ = 20 mV, *A* = 5 mV ms, and *D* = 0.74 (mV)^2^ms. **Left**: The triple-wise interaction computed from *simulated* spike sequences (gray lines: 50 individual trials each containing about 2500 spike occurrence of common input; dots and error bars: mean± standard deviation) compared with the *analytic* result of the mixture model (red line, Eq. 16) for *λ* = 5 Hz. **Right**: Analytical value of *θ*_123_ as a function of the bin size, Δ, for different common input rates, *λ*. **(B)** The triple-wise interaction when the postsynaptic neurons, blue ones, are in a *triangular architecture*. Each pair of postsynaptic neurons shares an independent common excitatory input. All parameters are as in (A). **(C)** Triple-wise interactions of 3 neurons receiving common excitatory or inhibitory inputs under star (top panel) and triangular (bottom panel) architectures. *θ*_123_ is represented as a function of *scaled diffusion coefficient*, 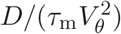, and *scaled shared signal strength, A/*(*τ*_m_*V*_*θ*_); the other parameters are: Δ = 5 ms and *λ* = 5 Hz. The white region in the bottom left panel shows the numerically indeterminate region due to a very small diffusion coefficient (level of noise) and strong inhibition. The right-hand side figures are for excitatory and the left-hand side ones are for common inhibitory inputs.

Figure 5C shows triple-wise interactions under star (Top) and triangular (Bottom) architectures for excitatory (right) and inhibitory (left) common input as a function of scaled diffusion coefficient (level of input noise) 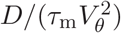 and scaled amplitude *A/*(*τ*_m_*V*_*θ*_). A single common excitatory input, in the star architecture, significantly increases the probability that all three neurons spike in the observation time window of Δ, *P* (1, 1, 1), whereas a single common inhibitory input increases the probability of the reverse pattern, *P* (0, 0, 0). This simply changes the sign of *θ*_123_ in Eq. 16. In the triangular architecture, with common excitatory input, however, each common input causes postsynaptic spikes for two neurons and does not drive the other one. This primarily increases *P* (1, 1, 0) (or any of its permutations) and attenuates the fraction (Eq. 16) that makes *θ*_123_ negative. For common inhibitory input, the probability of the reversed pattern, *P* (0, 0, 1) (or any permutations) increases; this results in a larger numerator in Eq. 16 and positive triple-wise interaction. These results demonstrate that not only the type of common input (excitation or inhibition) but also the underlying architecture (star or triangular) determines the sign of triple-wise interactions.

### Network structure and common input type can be determined from neural activity in vivo:Comparison with experimental data

The above observations raise a question: Is it possible to determine the type of common input and the underlying architecture from the event activity of neuronal population? Figure 6A shows the first-order parameter, 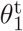, in a star or triangular architecture receiving either common excitatory or inhibitory inputs. The 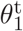 strongly depends on 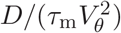, which measures the level of the noise in background inputs, but shows a weak dependence of the signal’s amplitude, *A/*(*τ*_m_*V*_*θ*_). More importantly, it does not show any conclusive dependence, neither on the choice of architectures nor type of common input. Thus, it is not possible to identify the underlying architecture nor the type of common input using the first-order parameter only.

**Fig 6.**
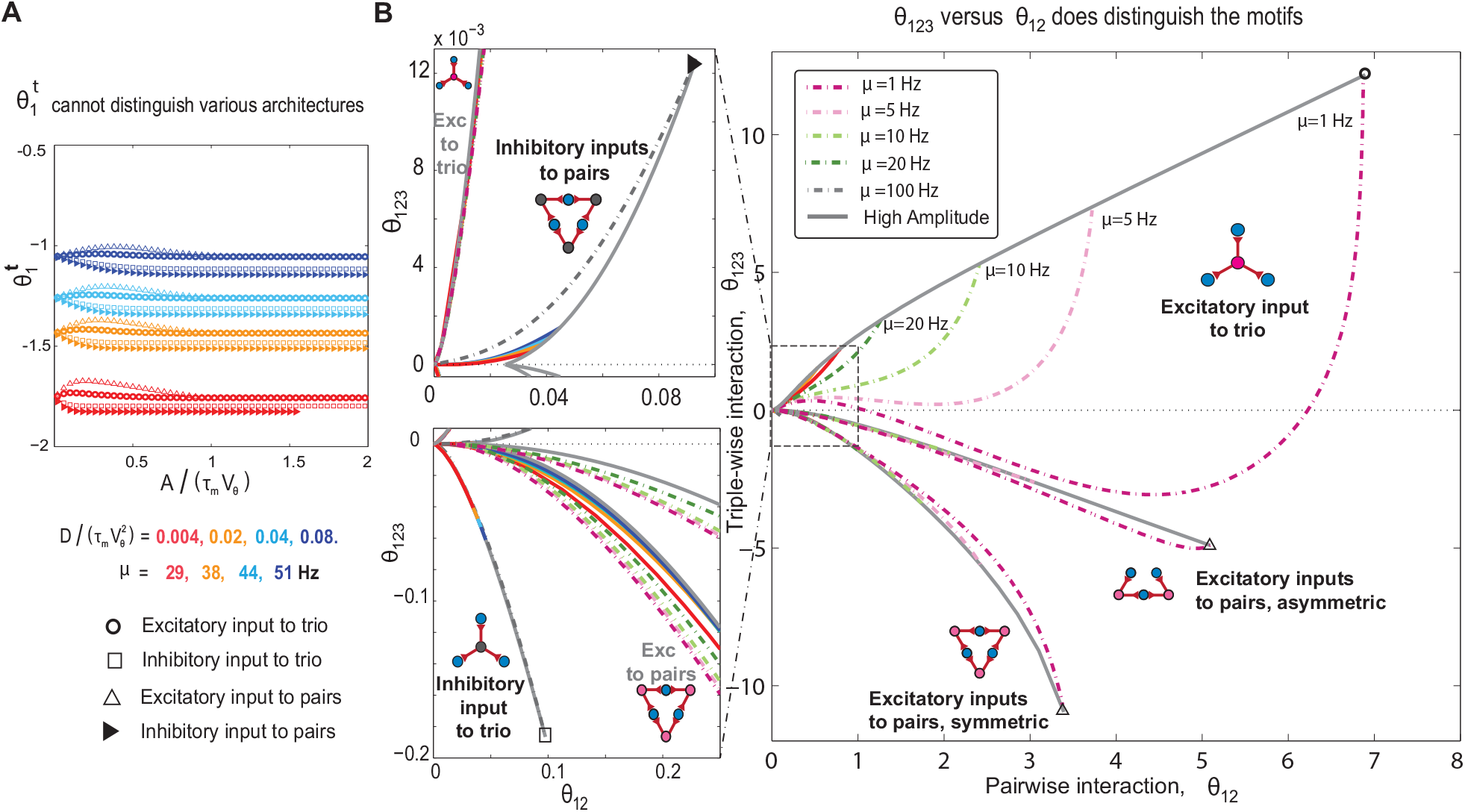
Population activity of three neurons caused by hidden common inputs occupies distinct regions in the plain of interactions, depending on the architecture and synaptic types. **(A)** The natural parameter for individual neurons, 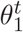, in *star* and *triangular* architectures with common *excitatory* or *inhibitory* inputs (four cases, shown by four symbols) versus *scaled* shared signal strength *A/*(*τ*_m_*V*_*θ*_). The 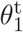 significantly varies with 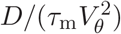, which measures *background spontaneous activity, µ*. However 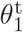 does not show any significant dependence on the shared signal strength, nor any conclusive dependence on the type of architecture. **(B)** In the plane of the triple-wise versus pairwise interactions, the star and triangular architectures with common excitatory or inhibitory inputs are distinguishable from each other. The regions associated with motifs of *excitatory common* input are expressly shown: From top to bottom, they attribute to *excitatory-to-trio, asymmetric excitatory-to-pairs*, and *symmetric excitatory-to-pairs* respectively. The regions associated with inhibitory common input are confined to around origin (i.e., small *θ*_123_ and *θ*_12_), hence are shown with higher resolution in left panels. Each region is bounded by two analytic *boundary lines*; the gray solid lines attribute to the high signal amplitude, i.e., |*A*|*/*(*τ*_m_*V*_*θ*_) ≫ 1, see SI; while the purple (gray) dashed boundary lines for motifs with excitatory (inhibitory) common input attribute to low (high) diffusion limit. The low (high) diffusion limit corresponds to background spontaneous activity of *µ* = 1 Hz (*µ* = 100 Hz), see SII. We choose these limits to cover a wide range of postsynaptic spontaneous activity while maintaining the assumption of low activity rates: *µ*Δ ≃ *F*_0_(Δ) ≤ 0.5, see SI. The colored solid lines are numerical results of the LIF neuron model for different scaled diffusions and signal amplitudes (Fig. S1). The fixed parameters are the bin size of Δ = 5 ms and the presynaptic rate of *λ* = 5 Hz.

Nonetheless, the 2D plane of *θ*_123_ versus *θ*_12_ does differentiate motifs, *c*.*f*. Fig. 6B; there, each motif clearly occupies its *distinct region*. Figure 6B shows 5 = 2 + 3 motifs in the *θ*_123_ versus *θ*_12_ plane. Two inhibitory motifs (*triangle* and *star*) occupy tiny areas; thus are expressly shown in two panels on its left (top and bottom). The three excitatory motifs cover much wider areas. Here we have also considered the *asymmetric excitatory-to-pairs*; it is the only other architecture with shared excitatory inputs, which can produce non-zero *θ*_123_. All five regions initiate from origin, i.e., *θ*_12_ = *θ*_123_ = 0; simply because both interactions vanish for zero signal’s amplitude, i.e., *A/*(*τ*_m_*V*_*θ*_) = 0. As we increase the signal’s strength, both *θ*_12_ and *θ*_123_ deviate from zero.

The example of excitatory-to-trio is the most visible case (SI, for comparable trends in other architectures). Consider postsynaptic neurons with the fixed spontaneous rate of *µ* = 1Hz (dashed-dotted purple curve); this curve shows how interactions change as we increase the shared signal’s amplitude from zero to highest conceivable values, i.e., *A/*(*τ*_m_*V*_*θ*_) ≫ 1. The pairwise interaction monotonically increases with the signal’s strength; but eventually saturates at its maximum of *θ*_12_ = 6.87, see the hollow black circle. The triple-wise interaction, however, shows a *non-linear behaviour*: It initially increases to *θ*_123_ = +0.34, then decreases to the negative value of −3.06, and finally increases to its saturation value of *θ*_123_ = +12.11, again the hollow black circle. We analytically show that for any choice of the spontaneous firing rate *µ*, we reach its corresponding saturation point, at the high enough signal’s strength (SI). The position of each saturation point (i.e. its *θ*_12_ & *θ*_123_) is found to be independent of the *neuron model*, as well as the *near the threshold* assumption. Conclusively, the saturation points (thick gray curve) form a *universal upper boundary* in the *θ*_123_ versus *θ*_12_ plane; the corresponding point for any *excitatory-to-trio* motif is placed below it (Fig. 6B). To address the *lower boundary*, we have limited ourselves to spontaneous rates *µ* ≥ 1Hz. As seen in Fig. 6B, for any higher value of the background activity (*µ >* 1Hz), the corresponding curve appears above the mentioned curve for *µ* = 1Hz, yet below the upper boundary of saturation points. Thus, practically, the curve for *µ* = 1Hz acts as the lower boundary.

A similar story holds for the other four motifs. Each corresponding region is composed of a bunch of curves. To obtain each curve, we assume a certain value of postsynaptic spontaneous rate, *µ*, then let the shared signal’s amplitude vary from zero to very high values. This produces a curve that initiates from the origin and ends at its saturation point. Each region is the accumulation of all these curves, and has two boundaries: One boundary is composed of all saturation points (thick-gray boundary), while the other is the curve with the lowest firing rate of *µ* = 1Hz (highest firing rate of *µ* = 100Hz), for motifs with excitatory (inhibitory) shared inputs, see Fig. 6B.

One significant question is how much the obtained boundaries vary with change in the *neuronal model* or the *near threshold regime* assumption. Can one region entirely displace, or even two *distinct regions* overlap? Fortunately, the high amplitude boundaries (thick gray curves) are *analytically* verified to be independent of the neuron model, as well as near threshold assumption (see SI). The other boundaries of low (high) spontaneous rate, for the excitatory (inhibitory) shared inputs, however, have a non-trivial behaviour. For the *star* architectures, they remain independent of neuron model; while for the *triangle* architecture, they do depend on the choice of neuron model, and the near threshold assumption (see SII). This dependence is actually an important example of non-linearity of input-output relation: It wouldn’t exist, if increasing the strength of presynaptic signal would linearly increase the probability of postsynaptic spike (technically, *F*_*A*_(Δ) = *F*_0_(Δ) + *cte* × *A*, see SII). However, this is only correct for *week signals*. In general, the probability of postsynaptic spike *non-linearly* varies with signal’s strength and saturates for very strong signals; the accurate description of this dependency requires a full knowledge of neuronal model (SII).

Figure 6 quantitatively answers the question we asked at the beginning of this part: It is possible to identify the underlying architecture and even the type of shared inputs (excitation or inhibition) for three homogeneous neurons, simply investigating their interaction parameters. As a practical example, we consider a careful study on V1 neurons of *anesthetized macaque monkey*; it investigated the relationship between the triple-wise interaction (Eq. 16) of three neurons (ordinate), and the average marginal pairwise interactions (Eq. 14) of neuron pairs in the group (abscissa) (Ohiorhenuan et al., 2010). They used extracellular recording of pyramidal neurons and found that many neurons, with mutual separations less than 300 *µm*, exhibited positive pairwise and strong negative triple-wise interactions (Fig. 7). The triple-wise interactions weaken as the electrodes’ separations increase above 600*µ*m. They attributed the observed *strong negative triple-wise* at near distances, to the hidden activity of small GABA-ergic inhibitory neurons, which presumably provide shared input to the larger excitatory pyramidal cells (Ohiorhenuan and Victor, 2011).

**Fig 7.**
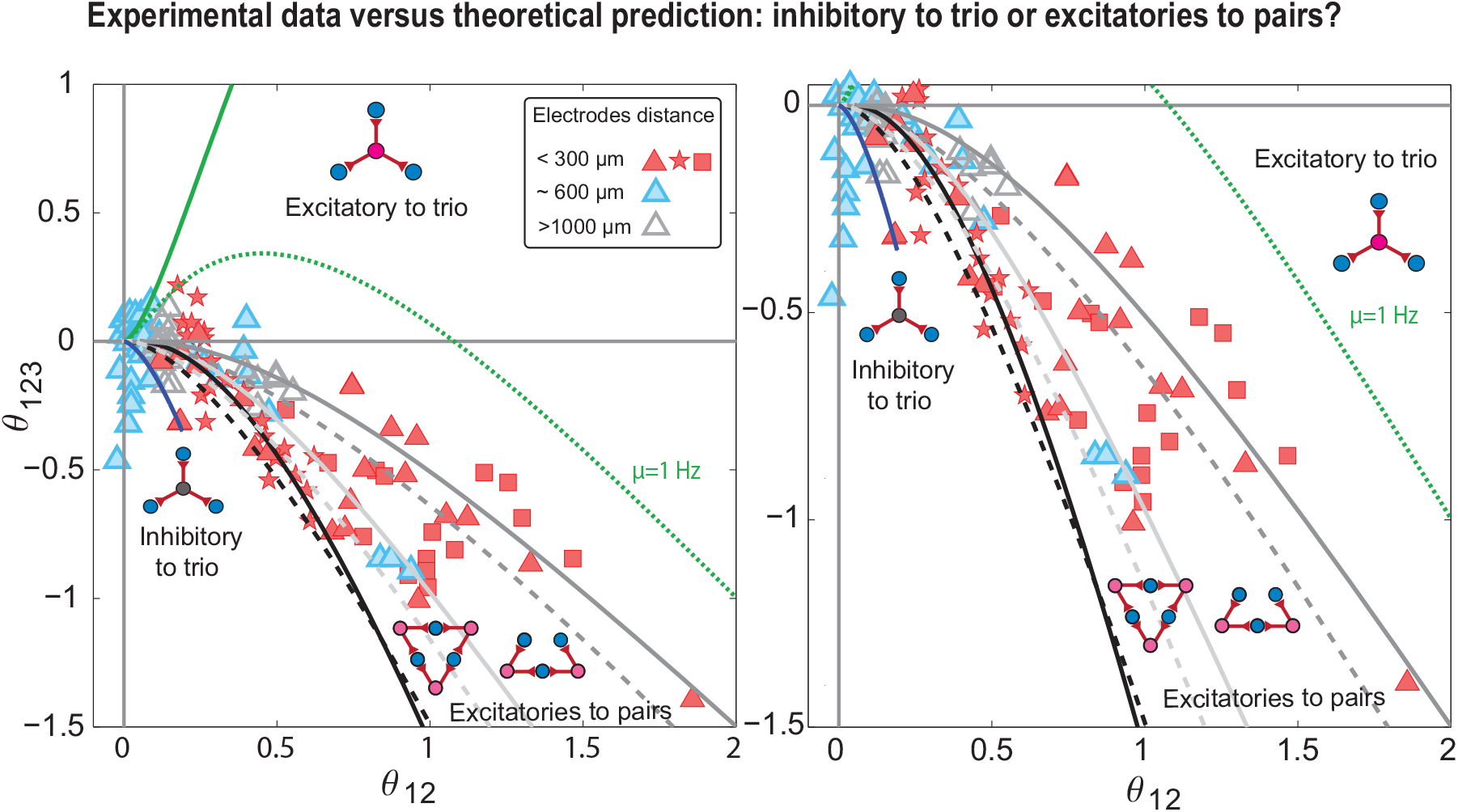
Comparison of theoretical predictions about neural interactions with experimental evidence extracted from (Ohiorhenuan and Victor 2011, Figure 4b). Ohiorhenuan et al. recorded the spike data from V1 neurons of macaque monkeys using tetrodes and analyzed the relation between triple-wise interaction (Eq. 16) of three neurons (ordinate) and an average marginal pairwise interaction (Eq. 14) of neuron pairs in the group (abscissa). Red and blue filled symbols represent the interactions of neurons within 300 and 600 *µ*m vicinity respectively while unfilled grey symbol shows the interaction at > 1000 microns distance. The black lines show the region of interactions theoretically obtained by assuming triangular architecture with common excitatory inputs to each pair of neurons. The solid black line is the high amplitude limit for interactions while the dashed black line is the boundary for the low spontaneous firing limit (*µ* = 1 Hz, low diffusion limit). Similarly, the dark gray solid and dashed lines determine the boundaries of region for triangular asymmetric architecture with common excitatory inputs given to two pairs among three neurons. The light gray lines are the same asymmetric excitatory inputs to pairs but all three pairwise interactions are taken into account for averaging while for the dark gray graphs, only nonzero pairwise interactions (for one pair, *θ*_12_ = 0) are considered. The blue analytic lines represent a narrow region for star architecture with common inhibitory input (high common input’s amplitude limit and high diffusion limit) whereas the orange lines show the wide region for excitatory inputs given to three neurons (dashed, low spontaneous activity limit, *µ* = 1 Hz for scaled diffusion limit of 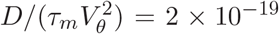 and solid, high common input’s amplitude limit). The fixed parameters are Δ = 10 ms and common input’s rate, *λ* = 5 Hz. The data reveals that nearby neurons (red filled symbols) receive pair of excitatory inputs in the triangular architecture rather than inhibitory or excitatory inputs in the star architecture.

Fortunately, the spontaneous activity of V1 neurons are reported within 10-70Hz (Gur et al., 1997; Ohiorhenuan et al., 2010; Ohiorhenuan and Victor, 2011), higher enough than the 1Hz lower boundary which we considered for the excitatory-to-trio motif. Thus, we can safely compare our theoretical predictions with the empirical observations. Figure 7 clearly shows that empirical data coincides with regions associated with the motif of excitatory-to-pairs, neither excitatory-to-trio nor any of the inhibitory motifs. It clearly rules out, the initial intuitive picture that shared inhibition induces the observed strong negative triple-wise interactions.

Inevitably, the following question emerges: Why should the observed strong negative triplewise interaction, be associated with excitatory common inputs, and the inhibitory shared inputs fail to produce any strong negative interactions? We will answer this question at the end of the results section. However, we firstly verify the robustness of our excitatory-to-pairs scenario, to the possible *complication of the motifs* due to recurrent interconnections, plausible change in *neuron model*, and even violation of *near threshold assumption*.

### Excitatory directional/recurrent connections among three neurons can explain the observed negative triple-wise interaction

Although the previous section’s analysis seems valid for common inputs’ architecture among three postsynaptic neurons, the question arises whether considering interconnections among the three neurons can induce any change to our concluding result in Figure 7. We, therefore, simulate all possible motifs that have directional or reciprocal connections, among the three neurons. At first glance, the number of motifs for three neurons is 2^6^ = 64 (each directed connection can be present or absent; so for 6 possible interconnections, it yields 2^6^). However, some of these motifs are structurally the same; they turn into each other, simply with changing (i.e. permuting) labels of three postsynaptic neurons. This means that 64 possible motifs could be further categorized in 16 main structures (Fig. 8). Regarding 16 main structures, we ask whether the inhibitory-to-trio with the help of either of them can reach the value of experimental data (red symbols in Fig. 7). Figure 8 shows the result of triple-wise interaction for each motif averaging over 50, 000 runs. We see four clusters of motifs, which are ordered as the number of inputs to pairs increases. The first cluster (blue, motifs 1 to 7) contains motifs with no simultaneous input from one excitatory neuron to two others (to pairs). For this cluster, the average triple-wise interaction is a small value. The second cluster (red, motifs 8 to 13) has motifs that have one excitatory neuron as the input to pairs of neurons in their architecture. The third and fourth clusters (green, motifs 14 and 15; and black, motif 16) contain architectures with respectively two and three excitatory inputs to pairs of neurons. Clearly, as the number of *excitatory-to-pairs* increases, the absolute value of triple-wise interaction boosts up, Fig. 8. The inset shows triple-wise versus pairwise interactions for these four clusters. This picture is consistent with our finding (shown in Fig. 7) that excitatory input to pairs induce large negative triple-wise and positive pairwise interactions. The excitatory input to pairs, either as common input or as directional connectivity, can generate such activity and thus is the basic architecture behind the data presented here.

**Fig 8.**
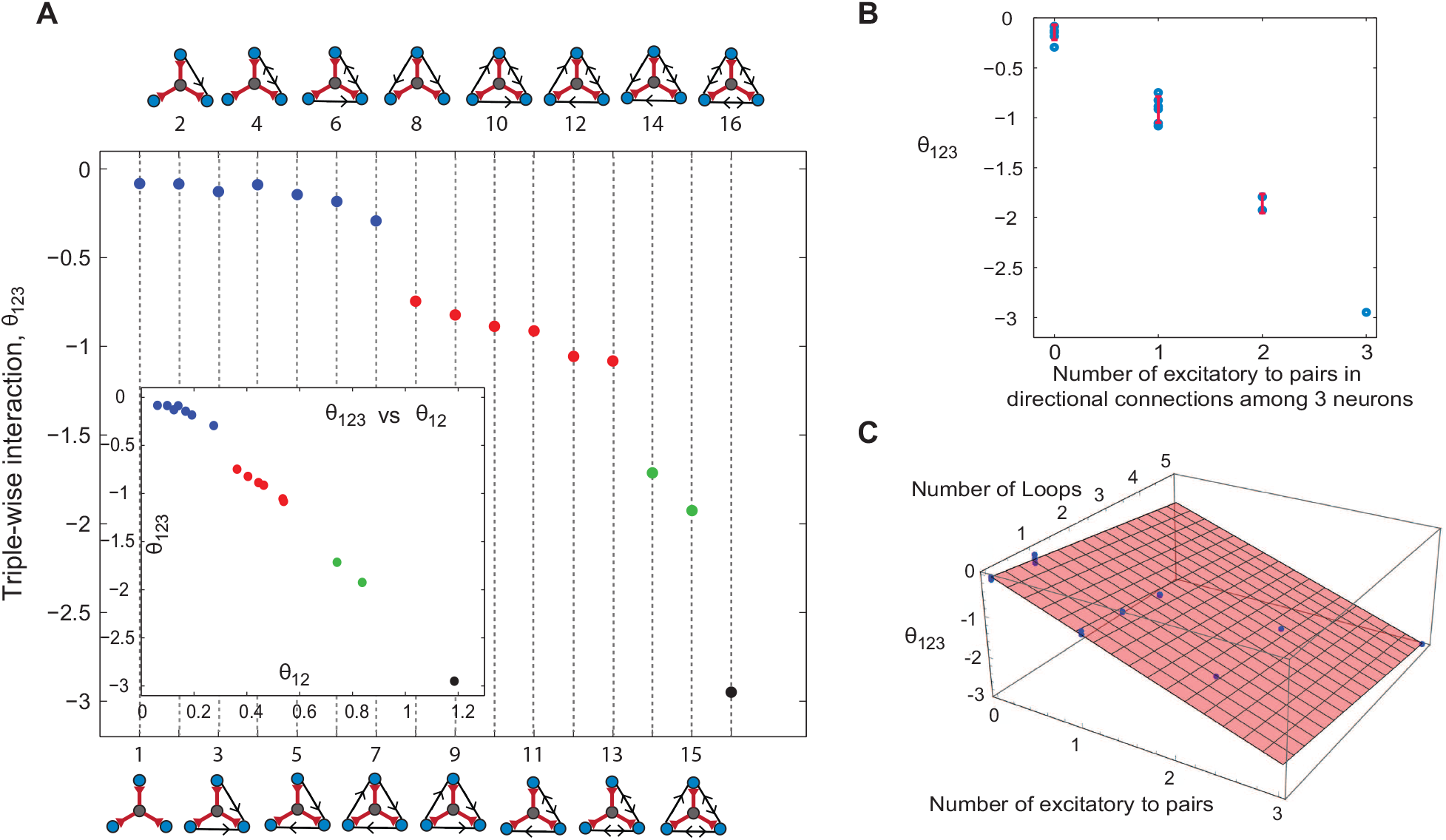
Triple-wise interactions for 16 motifs of directional and/or reciprocal connections among three neurons, when they receive independent noise and common inhibitory-to-trio. **(A)** The architectures are divided into four clusters based on the number of excitatory inputs to pairs in each motif. The first cluster (blue) contains directional connections that do not have any excitatory-to-pairs in their architectures. The second cluster (red) has one excitatory-to-pair in the motifs and the third and fourth (green and black) are related to two and three excitatory-to-pairs in their motifs. Inset shows the triple-wise interaction versus pairwise interaction for all 16 motifs. **(B)** The mean and error bar (standard deviation) of triple-wise interaction for each cluster as a number of excitatory-to-pairs. Clusters are separated from each other. **(C)** Triple-wise interaction for the 16 motifs is a linear function of both the number of excitatory-to-pairs motif and the number of loops. Each motif is simulated 50000 times, and each trial contains 500 seconds of spike trains with the time resolution of 0.05 ms. The time window to calculate the triple-wise and pairwise interactions is Δ = 5 ms, and the shape of presynaptic input is a square function for both common input and directional connection’s input, the same as analytic calculation. The parameters are: the scaled diffusion 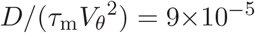, the scaled amplitude *A/*(*τ*_m_*V*_*θ*_) = 0.0125, the common presynaptic input and input rate of directional connectivity *λ* = 5 Hz, the time delay for directional interconnection *t*^′^ = 6 ms *>* Δ, and *A*^′^*/A* = 1, where *A*^′^ is the amplitude of the directional connections.

### Robustness: How does adaptation modulate the pairwise and triple-wise interactions?

It is a valid question, that how much our suggested framework (Fig.6), as well as its confrontation with the empirical data (Fig. 7), would change if we use a more physiologically plausible neuronal model? Regarding the predicted triple-wise and pairwise interactions, in Fig.6, we analytically prove that the curves for *high signal amplitude* are *independent* of the neuronal model, and the threshold assumption (see SI). This also holds for the curves with excitatory/inhibitory *to trio* architecture. Therefore, a good portion, not all, of the predictions would remain intact with or without a more plausible model. However, since a more physiologically plausible model can modify some predicted interactions for *excitatory-to-pairs* motifs; we have to study it, inevitably. The LIF neuron model is a good reduced model, that can reproduce in vivo spiking activity of neurons (Camera et al., 2004; Rauch et al., 2003). In comparison with data, however, it has some limitations and restrictions (Izhikevich, 2004; Jolivet et al., 2008; Ostojic and Brunel, 2011; Shinomoto et al., 1999). To have a more biologically plausible model, we add *adaptation term* to LIF (Brette and Gerstner, 2005; Gerstner et al., 2014a) and run simulation to see how the result of triple-wise versus pairwise interaction would change (more details about the model are in SIV). The simulation result shows adaptation reduces the firing rate of postsynaptic neuron (Fig. S4 and Fig. S5, SIV). As is explicitly shown at the end of the Results section, the lower background activity amplifies the effect of excitatory common inputs and diminishes that of inhibitory ones. This is why, the simulation results show that in the presence of adaptation, excitatory-to-pairs generates even stronger pairwise and triple-wise interactions, while inhibitory inputs induce weaker interactions (Fig. S6). Therefore, based on experimental evidence for low firing of V1 neurons (Ohiorhenuan et al., 2010), strong negative triple-wise interactions are induced by excitatory inputs to pairs motif, and *adaptation* simply strengthens this picture.

### Robustness: How do pairwise and triple-wise interactions change if we go slightly away from the threshold regime?

Alongside the problem of the plausible neuronal model, one can ask if the particular assumption of *near threshold regime* does reduce the scope of validity of our results. Fortunately, as mentioned earlier, many of the boundaries we have found are *independent of neuronal model*, hence independent of the near threshold assumption. Moreover, the particular near threshold assumption is partly asserted by an empirical study (Tan et al., 2014) that shows during sensory stimulation, V1 neurons operate *near the threshold regime* while background noise is uncorrelated.

Nevertheless, we run simulation to see how deviation from the threshold regime, could modify our results (see SIII). Comparing to the threshold regime, the simulations show that in the subthreshold regime, excitatory common inputs produce stronger interactions while inhibitory common inputs produce weaker ones. This trend simply reverses in the suprathreshold regime (Fig. S3, SIII). Now, considering the particular experimental study on macaque’s V1 (Ohiorhenuan et al., 2010), we recognize other evidence that cortical neurons operate in the *sub-threshold* (or near the threshold) regime (Shadlen and Newsome, 1998), depending on the state of the animal, and stimulus arrival (Tan et al., 2014). Assigning this fact to Fig.(7), it means that there is even a smaller region of inhibitory-to-trio which could be attributed to empirical data, due to the shift to subthreshold. While there is a larger portion of excitatory-to-pairs region which we can safely attribute to that data. This reaffirms our original conclusion that the observed strong negative triple-wise interactions are signature of excitatory-to-pairs, *exclusively*.

Finally, it is interesting to address why being in subthreshold (suprathreshold) results in increase (decrease) of the higher-order interactions induced by excitatory common inputs, and just do the reverse for interactions induced by inhibitory ones. Being in subthreshold (suprathreshold) mean lower (higher) spontaneous activity of postsynaptic neurons. As we show in the next part, it is the spontaneous activity that well explains all these observations.

### Excitation versus inhibition: Which one can produce stronger triple-wise interactions?

So far, we have found that the empirical strong negative *triple-wise* combined with positive *pairwise* interactions, for V1 neurons, are signature of microcircuits with *excitatory common inputs* (Fig. 7). One crucial question is why should other microcircuits with *inhibitory common inputs* have failed to produce such strong negative triple-wise interactions? Can we always attribute strong higher-order interactions to excitatory common inputs, or does it depend on certain features which vary from experiment to experiment?

The measured pairwise and triple-wise interactions depend on various features of the post-synaptic neurons, as well as their possible shared inputs. For the analytically tractable regime of *strong signaling inputs*, however, we can reduce many factors to a few decisive ones. Then, analytical calculations show that, when the spontaneous rate of postsynaptic neurons in time-window Δ is low, i.e., *F*_0_(Δ) ≪ 1, the excitatory common inputs can produce large pairwise and triple-wise interactions, while inhibitory common inputs can’t (Fig. 9, SI). This picture simply reverses if the spontaneous firing rate of postsynaptic neurons happen to be high, i.e., *F*_0_(Δ) ≲ 1. There is of course an intermediate regime, *F*_0_(Δ) ≃ 0.5, where the strength of interaction induced by *inhibitory input to trio* and *excitatory inputs to pairs* are nearly the same (Fig. 9).

**Fig 9.**
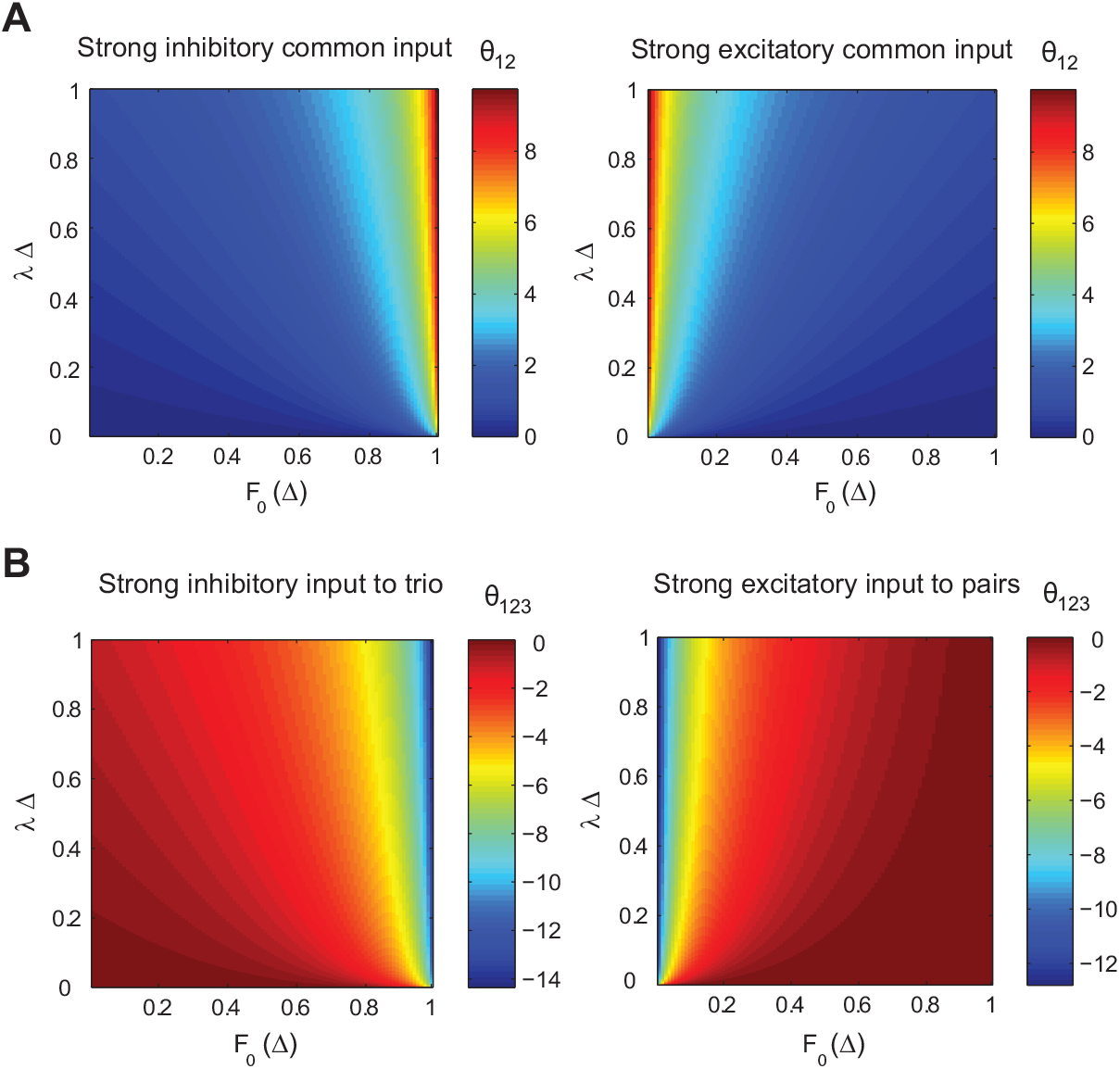
Pairwise and triple-wise interactions for strong excitatory and inhibitory inputs as a function of a spontaneous spike rate of a postsynaptic neuron (*F*_0_(Δ)) and an input rate (*λ*) in a small time window Δ. **(A)** inhibitory inputs generate strong pairwise interaction if the spontaneous rate of postsynaptic neuron is high (i.e., *F*_0_(Δ) ≃ 1) while excitatory inputs generate such strong interactions in a low spontaneous spiking regime, *F*_0_(Δ) ≪ 1. **(B)** The same story is true for triple-wise interaction independent of excitatory-to-pairs and inhibitory-to-trio architectures. Excitatory-to-pairs can generate strong triple-wise interactions in low spontaneous spiking but inhibitory-to-trio cannot induce such strong interaction in this regime.

Figure 10 illustrates how postsynaptic neurons’ spontaneous activity, i.e. *F*_0_(Δ), plays an essential role in relating the hidden underlying architecture with the observed interactions. If the regime of spontaneous rate is known, based on the statistics of neural data (pairwise and triple-wise interactions), one can predict the predominant architecture that induces the observed interactions. In a low spontaneous activity regime, motifs of excitatory inputs can induce strong triple-wise and pairwise interactions (regions in Fig. 10**A**); whereas, in high spontaneous activity, motifs with inhibitory inputs can generate strong interactions (Fig. 10**B**).

**Fig 10.**
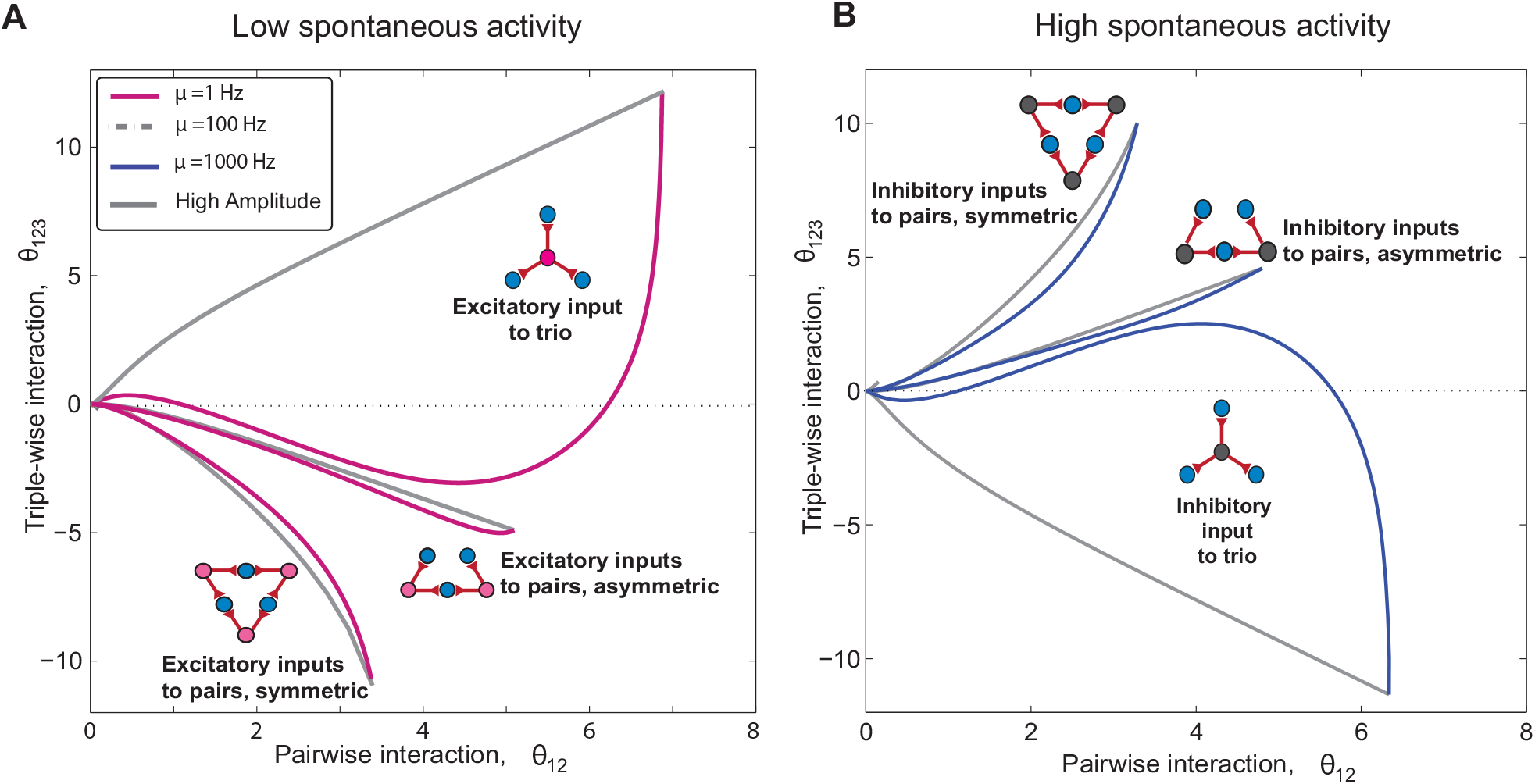
Uncovering the underlying architecture from observed higher-order interactions. **A** For spontaneous activity within the range *µ* = 1 −100 Hz, three regions for excitatory inputs’ motifs are shown. The strong higher-order interactions regardless of their signs are the sole signature of common excitatory inputs when the spontaneous rate is low. This scenario reverses if the background spontaneous activity is high (for example, olfactory bulb (Burton and Urban, 2015)). **B** The regions for spontaneous activity within the range of *µ* = 100 −1000 Hz show that inhibitory inputs can induce strong interactions in a high spontaneous regime. The excitatory inputs’ regions for high spontaneous activity (Fig. S2) shrink to a small size compared to the low spontaneous activity regime. The method for drawing the boundaries is similar to Fig. 6 and SII. The fixed parameters are bin size, Δ = 5 ms, and input rate, *λ* = 5 Hz.

In the experiment by Victor and colleagues (Ohiorhenuan et al., 2010; Ohiorhenuan and Victor, 2011) the neuronal firing rates ranged within 10Hz ≤ *µ* ≤ 70Hz (Fig.4 in (Ohiorhenuan and Victor, 2011)) while the time bin was Δ = 10ms. The exact spontaneous firing rate of postsynaptic neurons is *F*_0_ = 1 −exp(−*µ* × Δ); this yields 0.1 ≤ *F*_0_ ≤ 0.5. For such values of *F*_0_, any observation of strong triple-wise interactions is an indication of excitatory common inputs as opposed to inhibitory ones. Furthermore, there is an interesting fact in the data from Ohiorhenuan and Victor (Ohiorhenuan and Victor, 2011): The neurons that have the lower firing rate, generate stronger pairwise interactions; we particularly compare Fig.4c and 4a in (Ohiorhenuan and Victor, 2011), each set of data that has the smaller firing rate, produces larger pairwise interaction. This observed decrease of pairwise interaction with increasing the spontaneous activity (firing rate), is also revisited for *excitatory-to-pairs* architecture in figure 9-right, and not for the *inhibitory-to-trio*, see Fig. 9-left. This is another fact which reaffirms our original claim that the motif of excitatory inputs to pairs is the architecture behind this set of data.

## Discussion

### Overview

Our results point to the possibility of revealing the underlying neuronal architectures and the type of common input by using pairwise and triple-wise neural interactions (Fig. 10). Furthermore, for a specific set of empirical observations (Ohiorhenuan et al., 2010) in comparison with our analytical result, we show that excitatory shared inputs to pairs rather than intuitive inhibitory inputs to trio explain the data. Considering directional connections among three neurons does not disturb this picture: Excitatory inputs to pairs either as hidden layer common input or directional connectivity input are sufficient (and necessary) to explain the observed *strong negative triple-wise* and positive pairwise interactions. We also investigate the robustness of our result when the neuron’s voltage is *slightly away from threshold* (SIII) and when *adaptation* is present in the model (SIV). We analytically investigate how the *extreme case* of strong common inputs influence pairwise and triple-wise interactions (see SI); the analysis is *independent of any neuron model*, as well as *the voltage’s distance from the threshold*. It helps us to analytically find a *guide map*: Distinct regions with clear boundaries for each basic motifs in the triple-wise versus pairwise plane. This reveals that whenever spontaneous firing of neurons is *low*, motifs that have *excitatory inputs* can induce strong triple-wise interactions (Fig. 9); whereas when spontaneous firing rate is high, motifs with *inhibitory common input* can produce strong interactions. Likewise, a person among many others, if the majority are silent (low spontaneous activity), the one talkative person (excitatory input) is clearly noticed; whereas if the majority are talkative (high spontaneous activity), one silent person (inhibitory input) would be conspicuous.

### Comparison with other approaches

A classical approach to infer synaptic connectivity from extracellular spiking activity is to construct cross-correlograms of simultaneous spike trains from pairs of neurons Perkel et al. (1967). However, this approach aims at discovering connections among recorded neurons. In fact, researchers made efforts to eliminate the effect of common drives from unobserved inputs on this measure to avoid erroneously reporting pseudo-connections (Brody, 1999; Kobayashi et al., 2019). Another approach is a model-based method that uses a stochastic model of neurons. Among them, the point process - generalized linear model (GLM) is a standard tool for analyzing the statistical connectivity of observed neurons (Pillow et al., 2008; Truccolo et al., 2005; Volgushev et al., 2015). However, these models describe neuronal activity from their past activities and/or known covariate signals such as stimulus and local field potential signals. Since recorded neurons are embedded in larger networks, we need to take into account the effects of inputs from unobserved neurons in order to accurately describe the population activity. Although there have been attempts to include common inputs from unobserved neurons into the GLM framework by treating them as hidden variables (Kulkarni and Paninski, 2007; Vidne et al., 2012), variations in the structure of hidden common inputs are limited. In addition, these statistical models are not directly constrained by physiologically plausible membrane dynamics and spiking threshold while the LIF neuron model is (Ladenbauer et al., 2019). Here, given knowledge about the balanced network, we introduce hidden inputs as background noise and additionally consider various architectures of arbitrarily strong hidden common inputs as shared signals.

Another approach for modeling the input-output relation of a neural population under in-vivo conditions is to use the dichotomized Gaussian (DG) model (Amari et al., 2003; Macke et al., 2009, 2011) and its extensions (Montangie and Montani, 2015, 2017, 2018; Montani et al., 2013). Previous studies have shown that this simple model exhibits positive pairwise and negative triplewise interactions, which results in the observed sparse population activity (Shimazaki et al., 2015; Yu et al., 2011). The DG model is composed of threshold devices that receive inputs sampled from a correlated multivariate Gaussian distribution to model shared synaptic inputs (Amari et al., 2003; Leen and Shea-Brown, 2015; Macke et al., 2011). Limited by such a structure, one cannot test alternative hypotheses, e.g., if common inhibitory inputs can also generate the same neural interactions (Ohiorhenuan and Victor, 2011; Shimazaki et al., 2015).

In addition, the DG models do not incorporate the dynamics of synapses and membrane potentials. Conversely, the aforementioned input-output relation for near threshold neurons does address the dynamics of the membrane potential by using the LIF model neuron (Shomali et al., 2018); it thus yields quantitative results with temporal accuracy, enabling us to infer the types of common input under various architectures.

### Assumptions, limitations, and justifications of the framework

The quantitative model we introduced here is based on two distinct network architectures (triangle and star) with either excitatory or inhibitory shared inputs. It is crucial to see how the directional connection among postsynaptic neurons alters the view. The observed sparse connectivity of pyramidal neurons (Holmgren et al., 2003; Lefort et al., 2009; Markram et al., 1997; Mizusaki et al., 2016) shows that pyramidal neurons in visual cortex - in mature animals - are *not interconnected* (Jiang et al., 2015). However, the combination of directional connections with shared inputs is observed: Excitatory inputs from layer 4 are shared to layer 2/3 connected pairs of excitatory pyramidal neurons in cortex (Yoshimura et al., 2005). Hence, we ran simulations for the directional connections among three neurons: The results reaffirm that excitatory-to-pairs either in recurrent or common input’s motif, induces strong negative triple-wise and positive pairwise interactions in low spontaneous regimes (see Fig. 8). There is also another question, whether the *simultaneous existence* of common excitatory and inhibitory inputs in both triangle and star architectures damages this picture. We analyze the model in which two architectures are mixed, i.e., existing together and functioning simultaneously. The result shows the mixing of other motifs, while excluding excitatory-to-pairs, cannot induce strong negative triple-wise and positive pairwise interactions (see SV). It reiterates that the observed strong negative triple-wise interactions are the result of the *excitatory-to-pairs* motif. In other experimental results, of course, it is possible that the divergent common inhibition is mixed with the local common excitatory inputs. To extract evidence of the presence of such mixed inhibitory to trios from the data, one should carefully examine deviations from those observed interactions that are achieved solely by the excitatory inputs to pairs. As far as the spontaneous activities of neurons are low, however, we expect such deviations would be unfortunately small.

One of the assumptions for the analytical framework we introduced is that the firing rate of the signaling input is low (Wolfe et al., 2010) in comparison with postsynaptic neuron; so, there is at most one signal arrival during two successive spikes of the postsynaptic neuron. It is possible to consider cases with higher firing rates for signaling input (SIII in Shomali et al. 2018). However, as we consider a small time window, Δ = 10m*s*, the assumption of having *not more than one* signal arrival during such a short time window is practically acceptable. The other assumption is that the synaptic inputs set the voltage of the neuron near the threshold regime, which is reported to be the case when stimulus is presenting (Tan et al., 2014). We analytically calculate the *regions’ boundaries* for each motif (see Fig. 6): High amplitude boundaries for all motifs as well as low (high) diffusion boundary for excitatory-to-trio (inhibitory-to-trio) motif, are shown to be independent of the neuron model, hence the near threshold assumption. The other boundaries of triangle architecture, however, do depend on neuronal model, and the near threshold assumption. Yet, we carried numerical simulations, plus many other verifications, to make sure (i) the particular observed data on macaque V1 is signature of excitatory-to-pairs, and (ii) the suggested *guide map* practically remains reliable and intact, in more general situations. What does change the suggested *guide map*, and regions corresponding to motifs, are the spontaneous rate of neurons as well as bin size. Here, we compare the result for the infrequent and high spontaneous activity of postsynaptic neurons (Fig. 10) while assuming the low input rate, and small bin size. The bin size, on the other hand, cannot be too large, as it would diminish the effect of shared input (Fig. 5), hence the overall reliability of our formalism.

### Implication of the results and future challenges

Finally, we have attributed the observation of strong negative triple-wise interactions to a simple motif: Excitatory inputs to pairs. It is tempting to ask whether this microcircuit has any specified *computational advantage* so has been boldly observed, or whether it is the overall setting of a specific experiment that has resulted in this observation. On the one hand, there exist independent empirical evidence that the motif of excitatory-to-pairs is overexpressed - compared with a random network - in rat visual (Song et al., 2005) and somatosensory (Perin et al., 2011) cortex. In a recent theoretical study, this overexpression is explained as a consequence of maximizing capacity of the associative memory (Zhang et al., 2019); these observations suggest that the emergence of excitatory inputs to pairs is not an accidental observation. On the other hand, common inhibitory input has a clear computational advantage as a well-known winner-take-all network for sparse coding (de Almeida et al., 2009). There also exist experimental evidence that a common inhibitory input innervate multiple postsynaptic pyramidal neurons closer to each other than 100*µ*m (Packer and Yuste, 2011); at greater distances, the probability of common inhibitory inputs to two (and hence more) neurons decreases (Fig.6B in Packer and Yuste 2011). This is attributed to the limited length of inhibitory neurons’ axons and simply means that, if electrodes’ separation is greater than 100*µ*m, the chance of capturing a common inhibitory input (to pair, or to trio) has already diminished. For the particular experiment of Ohiorhenuan *et al*., the closest possible separation of recorded neurons is known to be less than 300 microns, *i*.*e*., recorded neurons are expected to be gathered in a circle of radius *r* ∼ 150*µ*m (Ohiorhenuan and Victor, 2011). Then, the probability of having neurons closer to each other than 100*µ*m is 32%, and that of having 3 neurons each closer than 100*µ*m to two others would be very low, less than 7% (see SVII). Thus, the probability of finding an inhibitory presynaptic neuron, which innervates synapses to three reocorded postsynaptic neurons was already less than 7%. Consequently, Ohiorhenuan *et al*. observation does not rule out the presence of inhibitory-to-trio architecture in a more local microcircuitry less than 100*µ*m, and cannot be used as empirical proof for the excitatory-to-pairs as an exclusive computational motif in microcircuits.

There need to be more precise experiments with higher control on the separation of electrode tips. If so, plotting how the observed triple-wise interaction varies with distances among neurons, would lead to a clear conclusion. Such a dependency for pairwise interactions varied with neurons’ distance is observed in retina ganglion cells (Ganmor et al., 2011). If the chance of observing a strong negative triple-wise interaction, for neurons closer than 100*µ*m, reduces, it indicates the absence of excitatory-to-pairs architecture in the local network less than 100*µ*m; therefore Ohiorhenuan *et al*.’s observation was a specific result of the experimental setting. However, if prevailing of strong negative triple-wise interactions persists even for neurons closer than 100*µ*m, it means that the excitatory-to-pairs are prevailing architecture in the microcircuit (≤ 300*µ*m), and would be another evidence to support for the *computational advantage* of *excitatory-to-pairs* microcircuit. On the contrary, it would be difficult to find evidence that inhibitory-to-trio exists or coexists with excitatory-to-pairs as computational units in the local microcircuits from activities of the three neurons as long as the postsynaptic firing rate is low, because of small negative values for triple-wise interactions induced by common inhibitory inputs (SI). Furthermore, although it is quite challenging to perform in vivo patch-clamp of common inputs and postsynaptic neurons at the same time, an experiment that can directly identify input types and the network’s structure in living animals is helpful to improve the prediction of this method. In summary, we have provided a theoretical tool based on the dynamics of a standard neuron model, to predict network architecture and types of hidden input neurons (excitatory/inhibitory) from the activity of neurons recorded in vivo. We define analytic regions for each motif, with boundaries mostly independent of neuron model, to show the basic motifs can be distinguishable from the statistical data. Our *guide map* helps to uncover hidden network motifs from neural interactions observed in a variety of in vivo data.

## Methods

### Leaky integrate-and-fire neuron at the threshold regime

According to (Shomali et al., 2018), the first-passage time density (inter-spike interval density) for the LIF neuron (Eq. 1) when neuron receives signaling input at time *τ*_*b*_ on top of noisy background input is given as

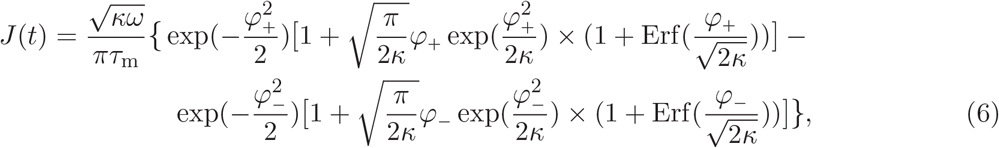

where 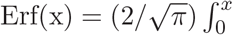 exp(−*t*^2^)*dt* and *κ*(*t, τ*_*b*_), *ω*(*tτ*_*b*_)and *φ*_±_(*t, τ*_*b*_) are:

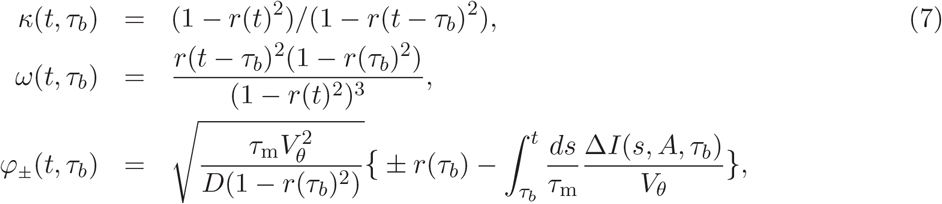

using *r*(*t*) = exp (−*t/τ*_m_). Before the occurrence of signal, i.e., *t < τ*_*b*_, the ISI density reduces to the known formula (Tuckwell, 1988; Wang and Uhlenbeck, 1945):

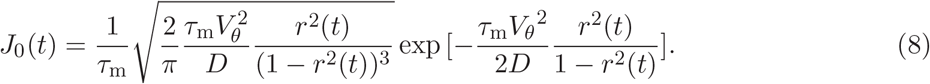

In this article, we use Eq. 6 with a square shape of signaling input given by:

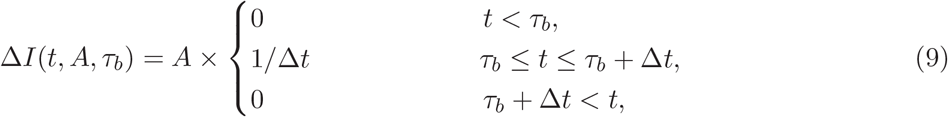

where Δ*t* measures the signal’s lasting time i.e., Δ*t* ∼ *τ*_s_, which is much smaller than *τ*_m_.

### Spiking density of LIF neuron after signaling input arrival

We derive the probability density of postsynaptic spike after the arrival of a signaling input. For this goal, we reset the *time origin* to signal arrival’s time. Following the aforementioned formalism in Eq. 6, the last postsynaptic spike would have happened at *τ*_*b*_ before the new origin. The conditional probability that the next postsynaptic spike happens at *τ* after signal arrival is calculated easily (Shomali et al., 2018):

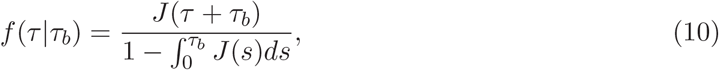

where the denominator is a normalization term to satisfy 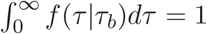. Next, we compute the probability density that the postsynaptic neuron has spiked at *τ*_*b*_ before signal arrival, but has not spiked since then, *p*_back_(*τ*_*b*_). It comes as the probability of Jbackward recurrence time, following renewal point process theory (Cox, 1962): 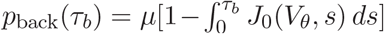, where 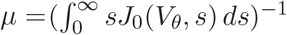 is the mean firing rate of the postsynaptic neuron when there is no signaling input. By marginalizing Eq. 10 with respect to *τ*_*b*_ using *p*_back_(*τ*_*b*_), we obtain (Shomali et al., 2018):

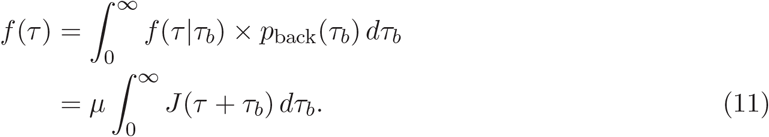

Note that when the amplitude of the signaling input, *A*, is reduced to zero, *J*(*V*_*θ*_, *t*) = *J*_0_(*V*_*θ*_, *t*) and *f* (*τ*) simplifies to 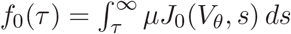.

Now, we can address the probability of having one or more spikes in a *specific time window* of Δ, after stimulus onset. It is given as the cumulative density function of *f* (*τ*):

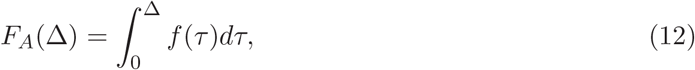

where subscript *A* indicates that *F*_*A*_(Δ) is a function of the amplitude of the signaling input. Δ Thus, in the absence of the signaling input (i.e., A=0) we have 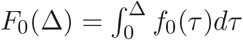.

### Pairwise and triple-wise interactions of neural populations

Using a binary representation of spiking activity for each postsynaptic neuron in a time window of Δ (schematically illustrated in Fig. 2), one can represent the population activity of the postsynaptic neurons as a binary pattern. From the probabilities of the occurrence of all possible patterns, one can assess pairwise or higher-order interactions of the neural population. For example, let us consider two neurons. Let *x*_*i*_ = {0, 1} (*i* = 1, 2) be a binary variable, where *x*_*i*_ = 1 means that the *i*th neuron emitted one or more spikes in the bin while *x*_*i*_ = 0 means that the neuron was silent.

We denote by *P* (*x*_1_, *x*_2_) the probability mass function of the binary activity patterns of the two postsynaptic neurons. Here *P* (1, 1) and *P* (0, 0) are the probabilities that both neurons are, respectively, active and silent within Δ. Similarly, *P* (1, 0) is the probability that neuron 1 emits one or more spikes while neuron 2 is silent during Δ; *P* (0, 1) represents the opposite situation. The probability mass function is represented in the form of an exponential family distribution:

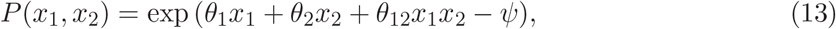

where (*θ*_1_, *θ*_2_, *and θ*_12_) are canonical parameters, and *Ψ* is a log-normalization parameter. In particular, *θ*_12_ is an information geometric measure of pairwise interaction (Amari, 2001, 2009b; Nakahara and Amari, 2002). Accordingly, the pairwise interaction parameter is computed:

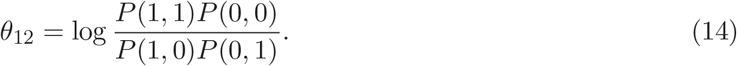

If the binary activities of two neurons are independent, *θ*_12_ = 0.

The same treatment is applied to three neurons. In an exponential form, the probability mass function for three neurons is written as

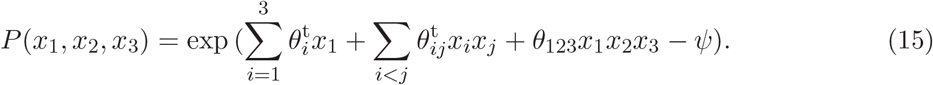

If *θ*_123_ (the triple-wise interaction parameter) is 0, the distribution reduces to the pairwise maximum entropy model, i.e., the least structured model that maximizes the entropy given that the event rates of individual neurons and joint event rates of two neurons are specified (Cover and Thomas, 1991). That is, a positive (negative) triple-wise interaction indicates that the three neurons generate synchronous events more (less) often than the chance coincidence expected from the event rates of individual neurons and their pairwise correlations. From this equation, the triple-wise interaction among three neurons for the exponential family of probability mass function is calculated using (Amari, 2009a; Nakahara and Amari, 2002):

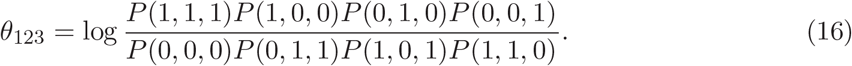

### Mixture model of three neurons receiving common inputs to their pairs (triangle architecture)

Here we explain the mixture model of three neurons whose pairs receive independent common inputs (a triangle architecture). As shown in Fig. 4D, there are 8 possible patterns to occur for the 3 independent common inputs. When the first common input is active (and the other two common inputs are silent), the pattern probabilities of three postsynaptic neurons are given by 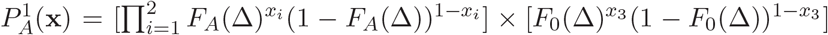. This situation happens in (*λ*Δ)(1 − *λ*Δ)^2^ × 100% of the bins. The probabilities of activity patterns in which neurons receive the second (third) common input, 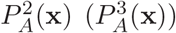, are given similarly to this equation. The common inputs may be simultaneously applied to the same bin due to their independence. Namely, two common inputs coincide at (*λ*Δ)^2^(1 −*λ*Δ) × 100% of the bins. The pattern probability in the bins at which common inputs 1 and 2 coincide is given by 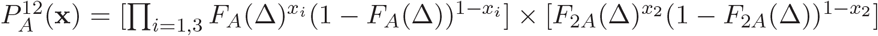. Similarly, we define 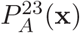 and 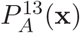 for the bins at which common inputs 2 and 3, and common inputs 1 and 3 coincide, respectively. Finally, all common inputs coincide at (*λ*Δ)^3^ × 100% of the bins, for which the pattern probability is given by 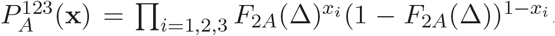. The parallel spike sequences are modeled as a mixture of these probability mass functions,

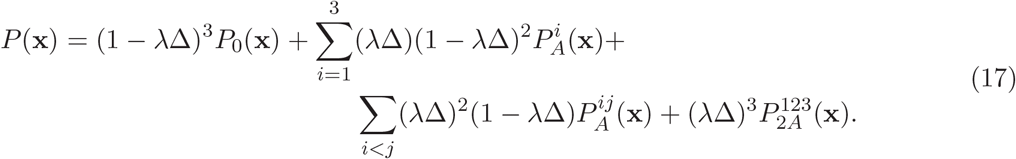

For the asymmetric case, when two common inputs are shared among three neurons (Fig. 4E), the mixture model simplifies to:

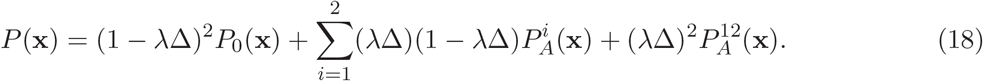

## Acknowledgments

S.R.S acknowledges J. Victor’s, and C. Clopath’s helpful comments; H.S. thanks S. Koyama and R. Kobayashi for the valuable discussions; S.R.S and S.N.R acknowledge T. Fukai’s kind support and are grateful to him and H. Maboudi for the valuable discussions. HS was supported by the Cooperative Intelligence Joint Research between Kyoto University and Honda Research Institute Japan, MEXT/JSPS KAKENHI Grant Number JP 20K11709, and the grant of Joint Research by the National Institutes of Natural Sciences (NINS Program No. 01112005).

## Supplementary materials

### SI: The pairwise and triple-wise interactions induced by strong inhibitory and common excitatory inputs

One main message of this paper is that the observed strong negative triple-wise interactions are signatures of excitatory-to-pairs (Fig. 7). The reasonable question is then why this should happen. Is there any fundamental difference between excitatory-to-pairs and inhibitory-to-trio such that the later one cannot produce strong negative triple-wise interactions? We approach this question by considering the extreme case of very strong signaling input. We firstly consider pairwise interactions and then extend the formalism to triple-wise interaction. Combining Eq. 14, Eq. 3, and Eq. 4, the pairwise interaction reads:

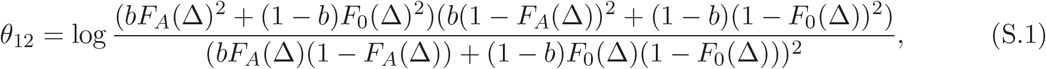

where *b* = *λ*Δ. If the postsynaptic neuron receives *an extremely strong inhibitory signal*, the chance of its spike in a short time-window of Δ immediately after signal arrival diminishes: *F*_*A*_(Δ) ≃ 0. This reduces Eq. S.1 to:

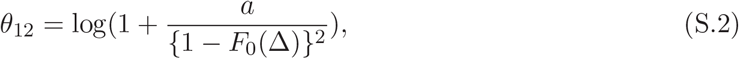

where *a* = *b/*(1 −*b*) = *λ*Δ*/*(1 −*λ*Δ). Conversely, for *an extremely strong excitatory signal*, the chance of spiking in the time window of Δ, immediately after signal arrival is almost 1; hence *F*_*A*_(Δ) ≃ 1. The pairwise interaction then simplifies to:

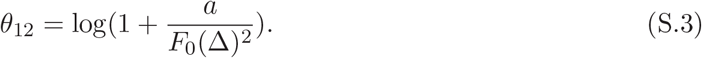

Comparing Eq. S.2 and Eq. S.3, we see that *F*_0_(Δ) is simply replaced by 1 −*F*_0_(Δ). In Eq. S.3, for excitatory signaling input and with *a >* 0, we obtain large *θ*_12_ when *F*_0_(Δ) ≪ 1. In fact, *θ*_12_ indefinitely increases as *F*_0_(Δ) → 0 ^+^. On the other hand, for strong inhibitory signals in Eq. S.2, this criteria changes to 1 −*F*_0_(Δ) ≪ 1; which means we need *F*_0_(Δ) ≲ 1 to obtain high *θ*_12_. Consequently, in a regime of low activity i.e., *F*_0_(Δ) ≪ 1, strong excitatory signals produce large *θ*_12_; whereas in high activity regime i.e., *F*_0_(Δ) ≲ 1, strong inhibitory inputs produce large pairwise interactions. The only approximation in the above reasoning is our assumption of low firing rate: *λ*Δ ≪ 1. More accurately, the Poissonian probability of signal arrival in a time window of Δ, is *b* = 1 −exp(*λ*Δ), which simplifies to *b* ≃ *λ*Δ in the low firing rate limit. Accordingly, the prefactor of *a* is, in general, *a* = {1 −exp(−*λ*Δ)}*/* exp(−*λ*Δ) which simplifies to *a* ≃ *λ*Δ*/*(1 −*λ*Δ) in the low firing rate limit. However, this correction does not disturb the overall result of our analysis at all.

Figure 9**A** shows how the pairwise interaction depends on spontaneous spiking rate and also the firing rate of signaling input in a small time window Δ for both inhibitory (left panel) and excitatory inputs (right panel). As the firing rate of signaling input increases, the amount of interaction induced by both excitatory and inhibitory inputs, increases as well. However, by increasing the spontaneous activity rate of *F*_0_(Δ), interaction induced by excitatory inputs to pairs decreases while that of inhibitory-to-pairs increases.

We extend this analysis to triple-wise interaction. Firstly, we consider the case of *strong inhibitory-to-trio*. Combining Eq. 15, Eq. 16, and Eq. 5 we have:

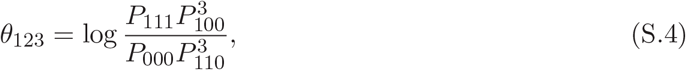

Where

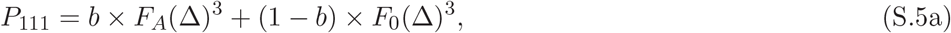

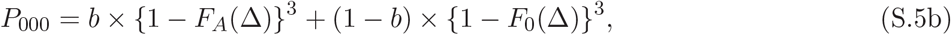

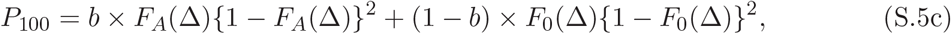

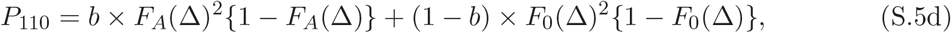

and *b* = 1 −exp(*λ*Δ), which simplifies to *b* ≃ *λ*Δ for low firing rates. Here, we have considered the aforementioned symmetry in the inhibitory-to-trio architecture, which enforces that *P*_011_ = *P*_101_ = *P*_110_, etc. In the limit of strong inhibition, for small time-window of Δ = 5 −10 ms, *F*_*A*_(Δ) ≃ 0. It drastically simplifies *θ*_123_ to:

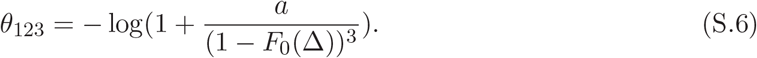

Here *a* = {1 −exp(−*λ*Δ)}*/* exp(−*λ*Δ), which is approximated to *a* = *λ*Δ*/*(1 −*λ*Δ) whenever *λ*Δ ≪ 1. For the case of *strong excitatory-to-trio*, triple-wise interaction becomes:

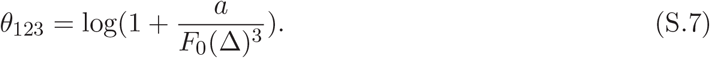

Similarly, we can consider the triple-wise interaction for excitatory-to-pairs architectures. It is a bit complicated compared to the inhibitory-to-trio case. Considering Eq. 15, Eq. 16 and Eq. 17, the triple-wise interaction is:

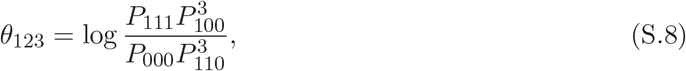

where

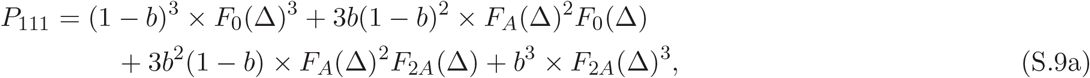

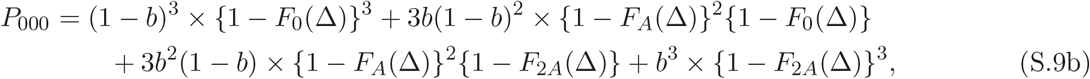

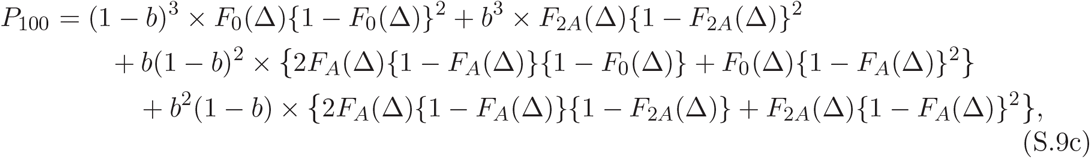

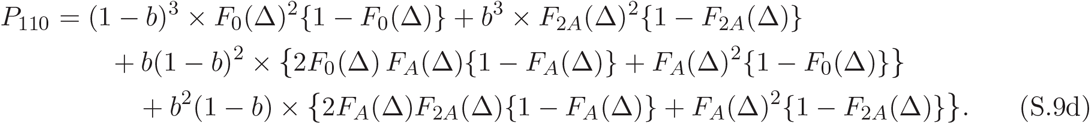

For extremely strong excitation to pairs, we have *F*_*A*_(Δ) ≃ 1 and *F*_2*A*_(Δ) ≃ 1; we thus have:

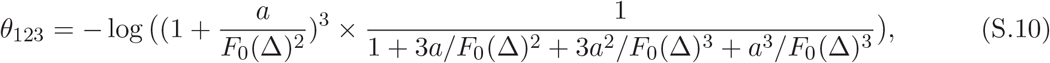

where *a* = {1 −exp(−*λ*Δ)}*/* exp(−*λ*Δ), as before. We can show that for the case of strong inhibitory inputs to pairs out of three neurons (*F*_*A*_(Δ) ≃ 0), the triple-wise interaction is:

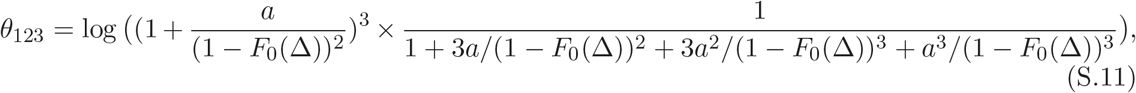

For asymmetric excitatory-to-pairs architecture (two common inputs instead of three), the pattern probabilities S.9a changes as:

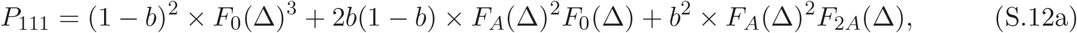

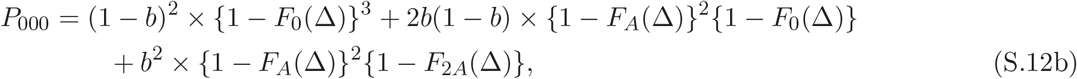

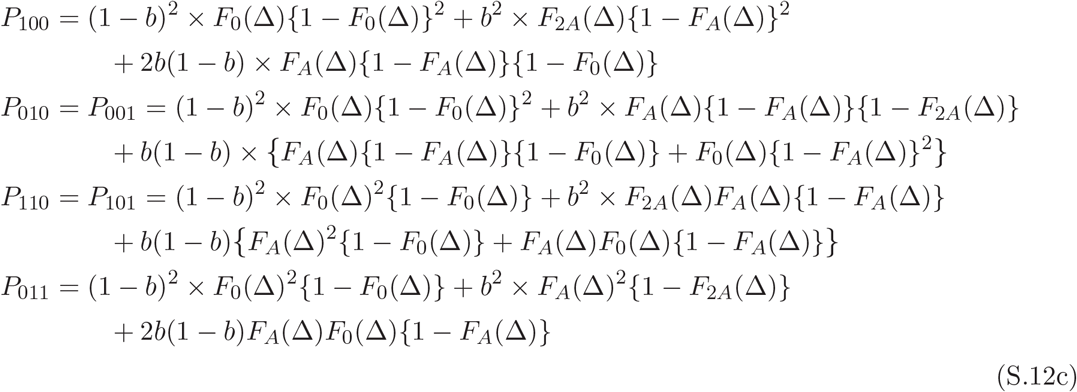

For a high amplitude limit of asymmetric excitatory inputs given to pairs of neurons (*F*_*A*_(Δ) ≃ 1), the triple-wise interaction becomes

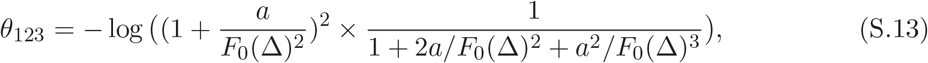

The marginalized *θ*_12_ for symmetric excitatory-to-pairs is (for inhibitory-to-pairs replace *F*_0_(Δ) with 1 −*F*_0_(Δ)):

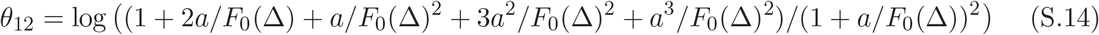

and for asymmetric excitatory-to-pairs is :

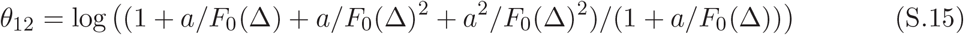

Figure 9B shows, for the *low activity* of a postsynaptic neuron, i.e., *F*_0_(Δ) ≪ 1, the motif of excitatory-to-pairs generates strong negative triple-wise interaction (right panel); whereas for *high spontaneous activity*, i.e., *F*_0_(Δ) ≃ 1, inhibitory-to-trio produces such strong results.

### SI-a: The limits of high amplitude approximation

In the high amplitude limit for excitatory inputs given to trio or pairs, the relations obtained for triple-wise and pairwise interactions (Eq. S.3 and Eq. S.7) in the limit of *F*_0_(Δ) → 1, diverge. For this regime, we take the limit of pairwise and triple-wise interactions (Eq. S.1, Eq. S.4 and Eq. S.5), considering *F*_*A*_(Δ) → 1 and *F*_0_(Δ) → 1 simultaneously. The pairwise equation, Eq. S.1, will be:

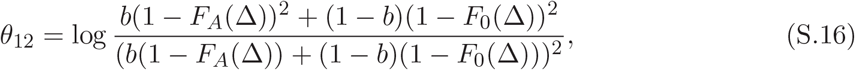

where *b* = *λ*Δ. Assuming *z* = (1 −*F*_*A*_(Δ)*/*(1 −*F*_0_(Δ)), it becomes

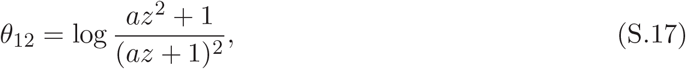

where *a* = *b/*(1 −*b*) and *z* varies within the range of [0, 1]. The triple-wise interaction (Eq. S.4 and Eq. S.5) in the limit of *F*_*A*_(Δ) → 1 and *F*_0_(Δ) → 1 is:

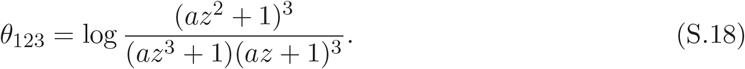

Similarly for inhibitory input given to three neurons, the limit is taken when *F*_*A*_(Δ) → 0 and *F*_0_(Δ) → 0 simultaneously. In this limit, we put 1 −*F*_*A*_(Δ) → 1 and 1 −*F*_0_(Δ) → 1. Then, pairwise interaction in Eq. S.1 reduces to:

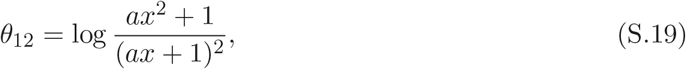

where *x* = *F*_*A*_(Δ)*/F*_0_(Δ). The triple-wise interactions (Eq. S.4 and Eq. S.5) will be:

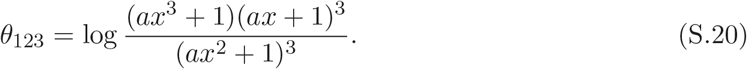

### SII: Defining the analytic regions for motifs of excitatory/inhibitory inputs to trios/pairs in the plane of *θ*_123_*/θ*_12_

Here we describe in brief how to achieve the boundaries for each motif in the plane of *θ*_123_*/θ*_12_. For all motifs, the first boundary arises from the limit of high amplitude (*F*_*A*_(Δ) → 1 for excitatory signaling inputs and *F*_*A*_(Δ) → 0 for inhibitory signaling inputs, see SI) when the spontaneous activity *F*_0_(Δ) changes within [0, 1]. This boundary is shown in Figure 6 by solid gray lines.

For the star architecture with common excitatory input, the second boundary (dashed purple line, Fig.6) is defined by the lowest plausible spontaneous firing rate of a postsynaptic neuron when CDF (*F*_*A*_(Δ)) varies from zero (no common input) to one (high amplitude of common input). One can easily find how spontaneous postsynaptic neuron’s firing rate relates to the level of background synaptic activities from a stationary distribution of voltage trajectory for LIF model neuron (Brunel, 2000; Brunel and Hakim, 1999). We have (SV in (Shomali et al., 2018)):

**Fig S1.**
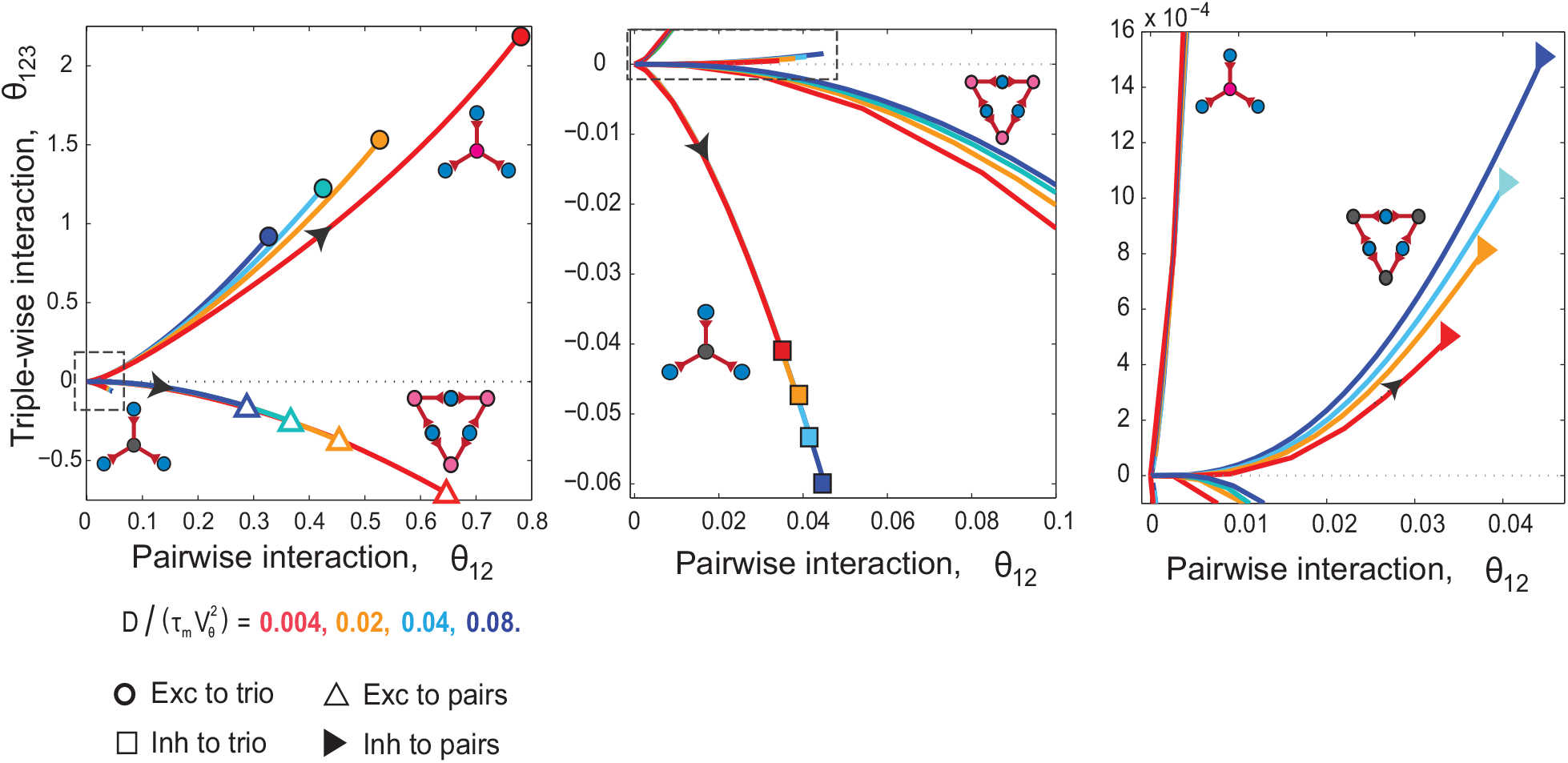
The triple-wise interaction versus pairwise interaction for star and triangular architectures with common excitatory and inhibitory inputs as a function of the scaled diffusion and scaled shared signal strength. The color codes are for four scaled diffusion coefficients and the symbols show the motifs. The arrows indicate increasing directions of the scaled amplitude parameter (i.e. *A/*(*τ*_m_*V*_*θ*_)) and the symbols on graphs show the saturation points. The middle and right panels illustrate the interactions in the neighborhood of origin for negative and positive triple-wise interactions which correspond to inhibitory-to-trio and inhibitory-to-pairs motifs, respectively. The only fixed parameters are the bin size of Δ = 5 ms and presynaptic rate of *λ* = 5 Hz.

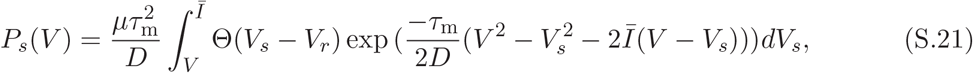

where Θ(*V*) is the Heaviside sJtep function: Θ(*V*) = 1 for *V >* 0 and otherwise Θ(*V*) = 0. *µ*is the mean firing rate: 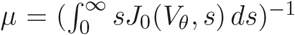 and *V*_*r*_ is the resting voltage that we assume to be zero here. *Ī* is the mean currentJwhich is equal to *V*_*θ*_ for the threshold regime. Since the stationary distribution is normalized 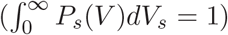, *µ* can be obtained explicitly (Brunel, 2000):

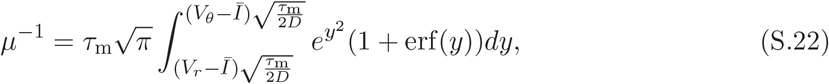

where 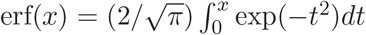. For *V*_*r*_ = 0 and *Ī* = *V*_*θ*_ (the threshold regime), the firing rate becomes

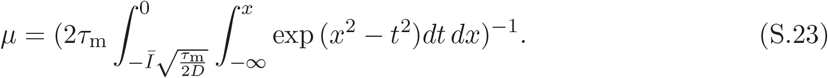

**Fig S2.**
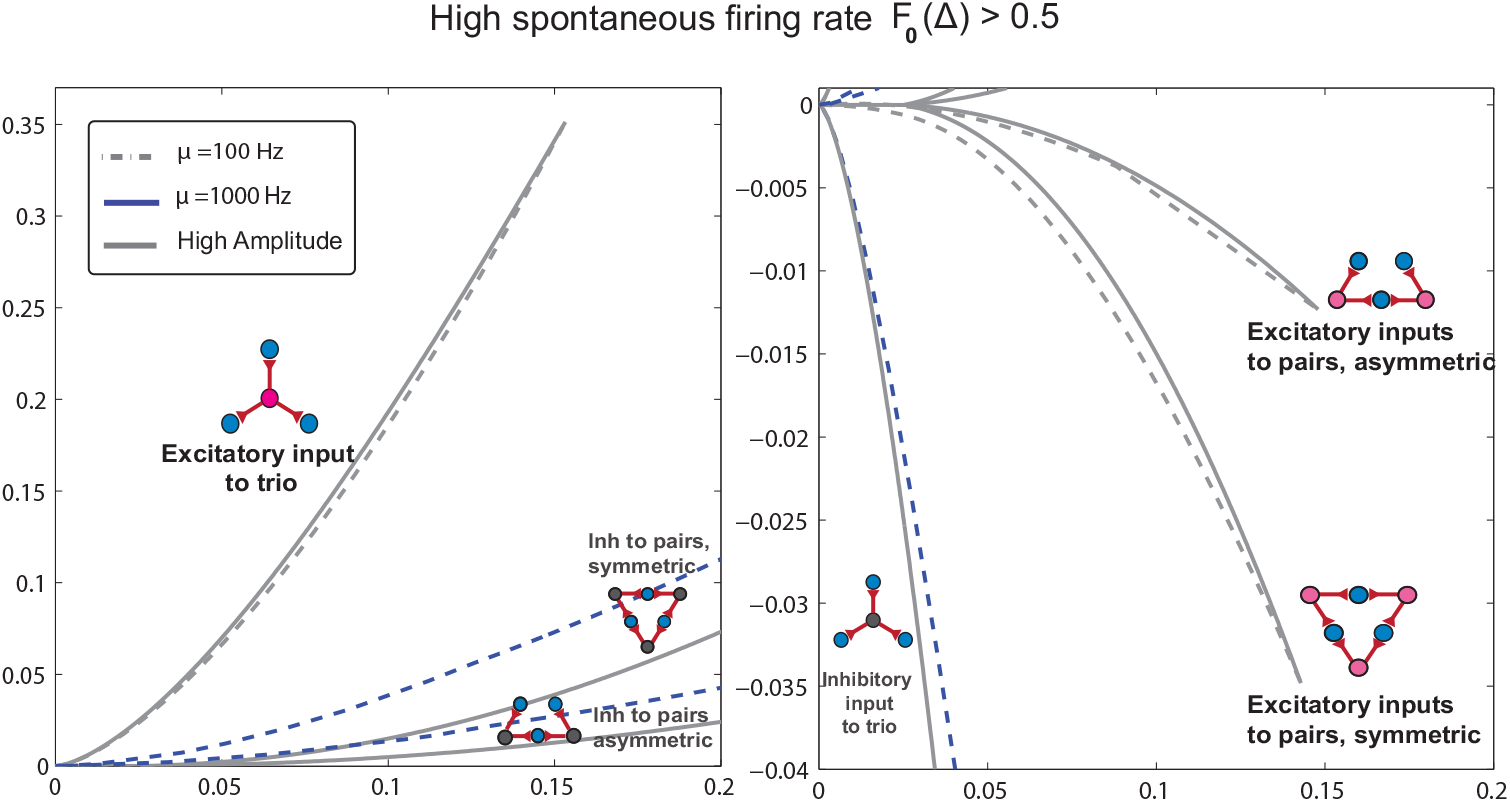
The regions of triple-wise interaction versus pairwise interaction for star and triangular architectures with common excitatory and inhibitory inputs for high spontaneous firing rates of postsynaptic neurons *µ* = 100 −1000 Hz (magnification of Fig. 10B). In this regime, the excitatory inputs’ motifs induce small interactions and their regions shrink compared to the low spontaneous firing of the postsynaptic neuron (Fig. 6). The only fixed parameters are the bin size of Δ = 5 ms and the presynaptic rate of *λ* = 5 Hz.

Using Eq. S.23, each level of background noise (diffusion coefficient) gives us a specific spontaneous firing rate that we can use to calculate *F*_0_(Δ). So by changing *F*_*A*_(Δ) within the valid range of [0, 1], we can calculate the whole range of triple-wise and pairwise interactions for excitatory inputs to trio (green to purple dashed lines in Fig.6). We observe that, as the background noise decreased, the region for excitatory-to-trio expands, so we use the lowest plausible spontaneous activity (here we use *µ* = 1*Hz* for *D* = 5.5 × 10^−17^*ms, mV* ^2^). This gives us the CDF of spontaneous activity, *F*_0_(Δ). By changing *F*_*A*_(Δ) within the range [0, 1], we obtain the second boundary for the motif of excitatory input to trio (Fig.6, purple line).

For inhibitory inputs given to trios, higher spontaneous activity shows higher interactions. So the second boundary is the limit of high spontaneous activity. Since we are restricted to a low activity regime (i.e., *F*_0_(Δ) ≤ 0.5), for this limit we choose the background noise for *F*_0_(Δ) = 0.5 that is (*D* = 190*ms, mV* ^2^ and *µ* = 100*Hz* for Δ = 5ms). Here, like excitatory-to-trio case, *F*_*A*_(Δ) varies within the range of [0, 1], but this time for high spontaneous activity (*µ* = 100 Hz). The two boundaries for inhibitory-to-trio, lie very close to each other and we get a narrow region in the plane of triple-wise/pairwise interactions.

For inhibitory inputs given to pairs of neurons, again the high spontaneous activity (here, (*D* = 190*ms, mV* ^2^ and *µ* = 100Hz) defines the second boundary (*F*_0_(Δ) = 0.5 for Δ = 5ms) which is achieved by our analytic input-output method and depends on the neuron model (Eq. 12, from Eq. 14 and Eq. 16).

Finally, the second boundary for the motif of excitatory-to-pairs in the low activity regime comes from very low diffusion (*µ* = 1*Hz* for *D* = 5.5 × 10^−17^*ms, mV* ^2^). The solution of postsynaptic neuron’s CDF after signal arrival (i.e. (*F*_*A*_(Δ) and *F*_2*A*_(Δ))) for very low diffusion regime is:

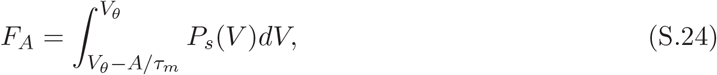

where *P*_*s*_(*v*) is the stationary distribution (Eq. S.21). When 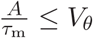, the CDF is:

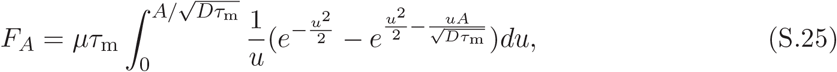

and for 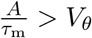:

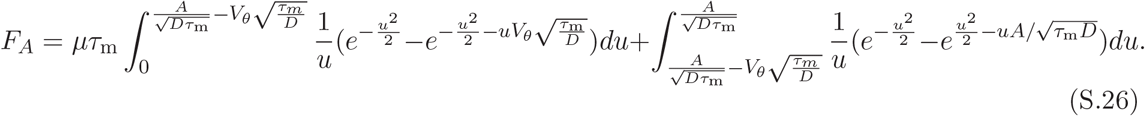

For a specific spontaneous firing rate of the postsynaptic neuron, from Eq. S.25 and Eq. S.26, pairwise and triple-wise interactions are calculated (Eq. 14 and Eq. 16) and drawn in lines in Fig.6. So the boundary is determined (*µ* = 1 Hz) for both symmetric and asymmetric excitatory inputs given to pairs of neurons (purple dashed lines in regions of excitatory inputs to pairs, Fig.6). In high spontaneous activity (Fig. 10B), the boundary for lowest spontaneous activity (*µ* = 100 Hz) of excitatory-to-pairs, is achieved by calculating the *F*_*A*_(Δ) and *F*_2*A*_(Δ) from input-output relation of a LIF neuron model in our method (Eq. 12) and then calculating interactions from Eq. 14 and Eq. 16. So just this boundary of excitatory-to-pairs like one boundary of inhibitory-to-pairs depends on the neuron model (here LIF).

### SII-a: When the motif of excitatory inputs to trio has negative triple-wise interaction?

We observe that when the spontaneous activity of the postsynaptic neuron is less than *µ* = 5 Hz, the triple-wise interaction in the motif of excitatory inputs given to three neurons, for some values of common input’s amplitude, becomes negative (Fig. 6). We can rewrite the pattern probabilities and triple-wise interaction for excitatory inputs given to three neurons from Eq. S.4 and Eq. S.5:

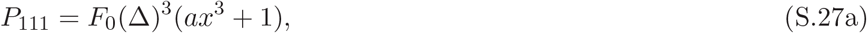

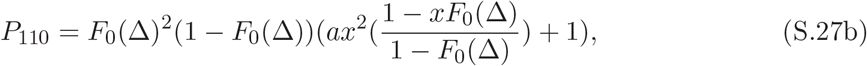

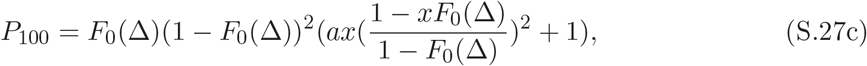

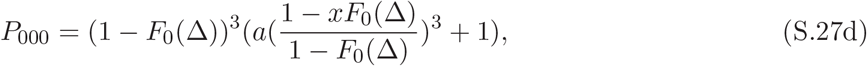

And triple-wise interaction is:

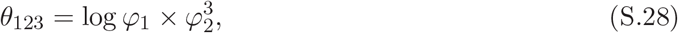

that

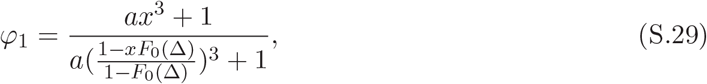

and

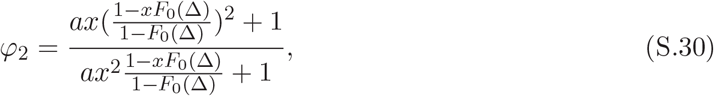

where *a* = *b/*(1 −*b*) and *x* = *F*_*A*_(Δ)*/F*_0_(Δ). For excitatory inputs to trio, *x* varies within the range 1 ≤ *x* ≤ 1*/F*_0_(Δ). The term that makes the triple-wise negative in some values of common input’s amplitude, is *ϕ*_2_, which is the fraction of pattern probability *P*_100_ over *P*_110_. By investigation, we find that when *η <* 0.25 where *η* = *F*_0_(Δ)(1 −*F*_0_(Δ))*/a*, the triple-wise interaction becomes negative in motif of excitatory-to-trio. For this condition, negative triple-wise interaction occurs when the cumulative density function *F*_A_ (Δ) is in the range of *F*_−_(Δ) ≤ *F*_*A*_(Δ) ≤ *F*_+_(Δ), where 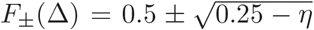 For example when the common input rate is *λ* = 5 Hz and time window is Δ = 5 ms, postsynaptic spontaneous rate below *µ* = 4.4 Hz, satisfy *η <* 0.25 and the triple-wise interaction becomes negative.

### SIII: What happens to triple-wise and pairwise interactions when we go beyond threshold regime?

Here we investigate how the results change when we go beyond the threshold regime. We performed simulations when the postsynaptic neurons are in the subthreshold and suprathreshold regimes, and compare the results with simulations at the threshold regime. Figure S3 shows the interactions arisen from the simulations in the plane of pairwise versus triple-wise interaction parameters. The circles and rectangles marked in orange display the results when the postsynaptic neurons operate at the threshold regime. With the same parameters, those marked in yellow and pink are obtained when the mean inputs to postsynaptic neurons reach 2 and 10 percent below the threshold (*δI/Ī* = −0.02, −0.1). The circles show the interactions obtained for the inhibitory-to-trio motifs, whereas the rectangles are for the excitatory-to-pairs motifs. The result is for three scaled amplitudes of the signaling inputs at *A/τ*_*m*_*V*_*θ*_ = 0.2, 0.4, 1 shown with no line, one line, and two crossed lines inside the symbols respectively. These results show that, as the mean input goes away from the threshold in the subthreshold regime, the interactions for excitatory-to-pairs motifs get stronger while those for inhibitory-to-trio get weaker. These results are interpreted as follows. At the threshold regime, the mean input to a postsynaptic neuron is set very close to the threshold. Thus, even a small amount of noise induces spikes of the postsynaptic neuron. In subthreshold regime where the mean input given to the postsynaptic neuron is set below the threshold, the neuron generates a spike as far as the variance of the noise helps the voltage to reach the threshold. In this case, the postsynaptic neuron fires less frequently (smaller amount of *F*_0_(Δ)) compared to the case at the threshold regime. Since the neuron is in a regime of lower spontaneous activity, the interactions induced by the excitatory inputs increase while the interactions caused by inhibitory inputs decrease (see SI). Therefore, the hypothesis that the excitatory-to-pairs motif is behind the empirically observed strong triple-wise and pairwise interactions is not only unchanged but also more strongly supported when the neuron operates in the subthreshold regime rather than the threshold regime.

**Fig S3.**
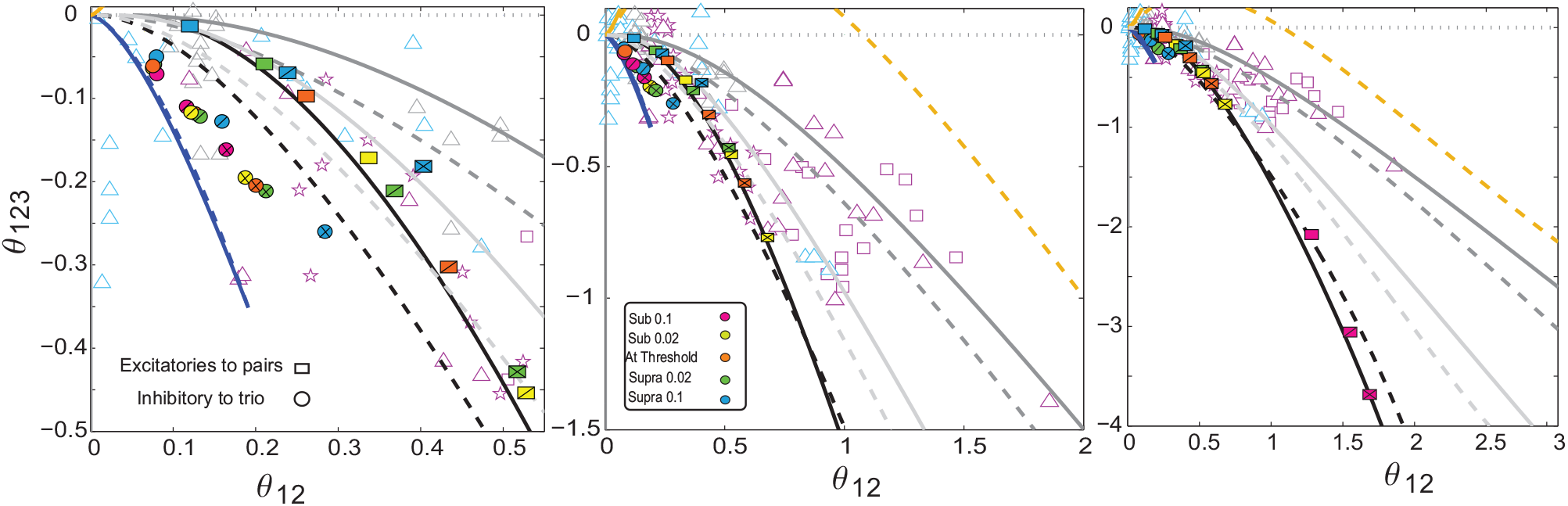
Modulation of interaction parameters in subthreshold and suprathreshold regimes, where mean synaptic input given to membrane potential is set below or above threshold. Left: Changes in interactions among inhibitory-to-trios are shown in subthreshold and suprathreshold regimes. Each color is for one deviation (see the color code) from the threshold regime at three scaled amplitudes of the signaling inputs at *A/τ*_*m*_*V*_*θ*_ = 0.2, 0.4, 1 shown by no line, one line and two crossed lines inside symbols respectively. Middle and Right: The interactions are shown for the motif of excitatory inputs given to pairs of neurons. When a neuron is in the subthreshold regime, the interactions are reduced for inhibitory-to-trio of neurons (compare orange with pink and yellow circles) while they are increased for excitatory-to-pairs motifs (compare orange with pink and yellow rectangles). In the suprathreshold regime, the excitatory-to-pairs motif generates weaker interactions (green and blue rectangles) while the inhibitory-to-trio motif generates stronger interactions especially for stronger input (green and blue circles with crossed lines inside). Fixed parameters are Δ = 10 ms, 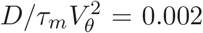, and *λ* = 5 Hz. Each symbol results from at least 10^10^ steps of run to keep the mean squared error in order of 10^−3^ −10^−4^.

We then move on to the suprathreshold regime, where the mean input is set above the threshold of postsynaptic neuron’s voltage (Figure S3). Those marked in green and blue are obtained when the mean inputs to postsynaptic neurons reach 2 and 10 percent above the threshold (*δI/Ī* = 0.02, 0.1). In the suprathreshold regime, the picture mentioned above reverses. Since the mean input is set above the threshold, a neuron generates regular spikes with high frequency, changing the regime of spontaneous activity from low to high firing rates *F*_0_(Δ) *>* 0.5. In this regime, the inhibitory signaling input can induce stronger interactions while excitatory signaling input cannot. We can reconfirm this interpretation in the simulation study (Fig. S3). For the range of scaling amplitudes of signaling inputs *A/τ*_*m*_*V*_*θ*_ = 0.2, 0.4, 1, we observe the interactions induced by inhibitory inputs get stronger as the mean input increases to 2 and 10 percent above the threshold (compare orange circle with the green and blue circle for symbols with crossed lines, for example).

### SIV: Does adaptation render inhibitory-to-trio to induce strong negative triplewise and positive pairwise interaction?

Here we want to use a more physiologically plausible model of the neuron and see how it affects the interaction parameters. For this aim, we add adaptation effect to the leaky integrate-and-fire neuron model (Brette and Gerstner, 2005; Gerstner et al., 2014b):

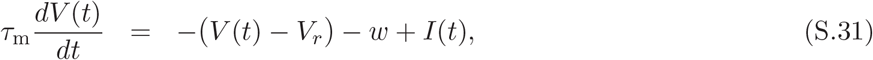

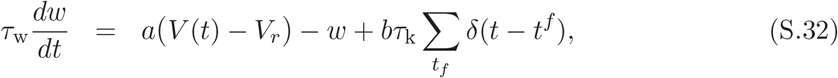

where the adaptation current is *w*, the coupling term of voltage to adaptation is *a*, and *τ*_w_ is the time constant of the adaptation variable. When the neuron generates a spike, the voltage is reset to *V*_*r*_ and the adaptation variable *w* is increased by *b* which is a parameter for the spike-triggered adaptation. We use adaptation parameters consistent with Brette and Gerstner (Brette and Gerstner, 2005). Figure S4 shows the result with and without the adaptation at the threshold regime for the two motifs of excitatory-to-pairs (rectangles) and inhibitory-to-trio (circles) for three scaling amplitudes of signaling inputs *A/τ*_*m*_*V*_*θ*_ = 0.05, 0.1, 0.25 shown with no line, one line and two crossed lines inside the symbols respectively. For adaptation, we consider two cases in that the coupling between voltage and adaptation current (*a*) is present or absent. The former (i.e. *a >* 0) considers subthreshold adaptation in addition to spike-triggered adaptation (*b*) while the latter (i.e. *a* = 0) assumes just the effect of spike-frequency adaptation. Figure S4 shows how adaptation modulates the pairwise and triple-wise interactions for inhibitory-to-trio and excitatory-to-pairs motifs. We observe that the modulation by adaptation make the triple-wise (pairwise) interactions for the excitatory-to-pairs motif more negative (positive), while the triple-wise (pairwise) interactions in the inhibitory-to-trio motif get less negative (positive) for strong inhibitory signaling inputs. It makes a larger difference in triple-wise and pairwise interactions between these two motifs.

The result shows adding the effect of adaptation differentiates, even more, the triple-wise interactions between the excitatory-to-pairs and inhibitory-to-trio motifs in a way that the excitatory input to pairs can induce the strong negative triple-wise and positive pairwise interactions, while the strength of inhibitory-to-trio’s interactions for strong common input decreases. This is attributed to decreasing the spontaneous activity of postsynaptic neurons in the presence of adaptation (Fig. S5). We observed that adding the effect of adaptation renders the firing rate of the postsynaptic neuron receiving common inputs on top of noisy background input, from 22 Hz (excitatory-to-pairs motif) and 19 Hz (inhibitory-to trio motif) to around 5 Hz and 3 Hz respectively (also Fig. S6, middle and right panels). When common input is off, adaptation also decreases the firing rate of postsynaptic neuron from 22 Hz to around 3 Hz (compare number 1 blue with red in Fig. S6, middle). Therefore by reducing the spontaneous activity, adaptation increases the strength of interactions in excitatory-to-pairs motifs but decreases the interactions’ strength in inhibitory-to-trio motifs (Fig. S6, middle and right panels).

**Fig S4.**
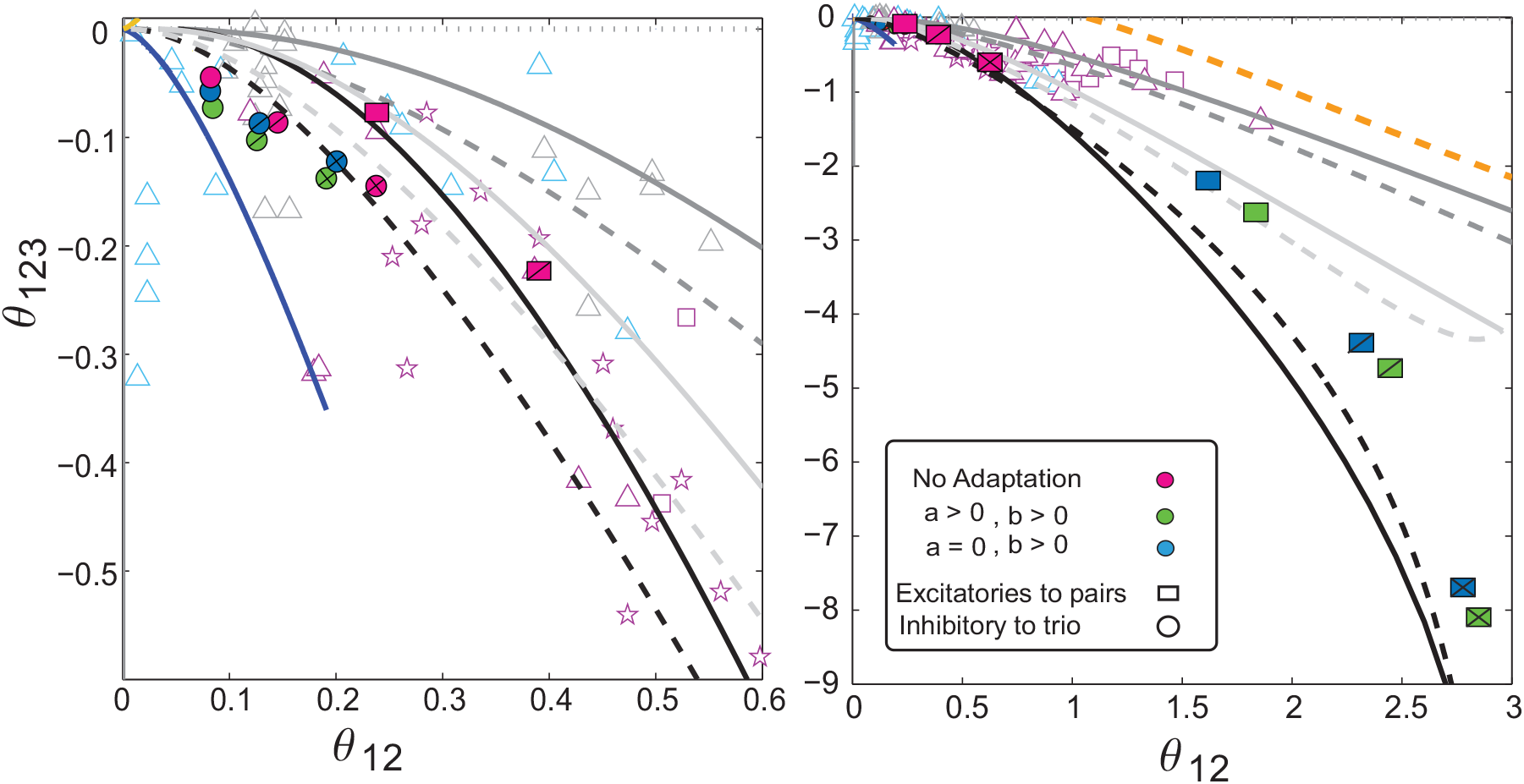
Modulation of interaction parameters in the presence of adaptation. The left panel shows the changes of interactions among the inhibitory inputs given to trios in the presence of adaptation (blue and green circles) compared with no adaptation case (purple circles). Each case is shown for three scaled amplitudes of the signaling inputs at *A/τ*_*m*_*V*_*θ*_ = 0.05, 0.1, 0.25 shown with no line, one line and two crossed lines, inside the symbols respectively; the larger the amplitudes, the stronger the interactions. When the adaptation is involved, the interactions are weaker for strong inhibitory input given to three postsynaptic neurons (left: blue and green circles) while they are stronger for excitatory-to-pairs motifs (right: blue and green rectangles). The adaptation parameters are consistent with Brette and Gerstner paper (Brette and Gerstner, 2005). Fixed parameters are *a* = 4 *nS* (for *a >* 0 cases), *b* = 0.0805 *nA, τ*_*m*_ = 10 ms, *g*_*L*_ = 30 *nS, C* = 281 *pF, V*_*r*_ = − 70 *mV, V*_*θ*_ = − 50 *mV, τ*_*w*_ = 144 ms, Δ = 10 ms, *D* = 0.74 *msmV* ^2^, and *λ* = 5 Hz. Each symbol results from at least 10^10^ steps of run to keep the mean squared error in order of 10^−4^.

The other interesting effect we observe is that excitatory inputs given to pairs of neurons with the help of adaptation can generate the observed CV (coefficient of variation) in V1 neurons (Gur et al., 1997), and by increasing the amplitude of signaling input, the CV reaches high values (Fig. S6 A) consistent with some experimental studies reporting high variability for neurons (Shadlen and Newsome, 1998). On the other side, the motif of inhibitory-to-trio cannot generate such a result of high CV (Fig. S6).

**Fig S5.**
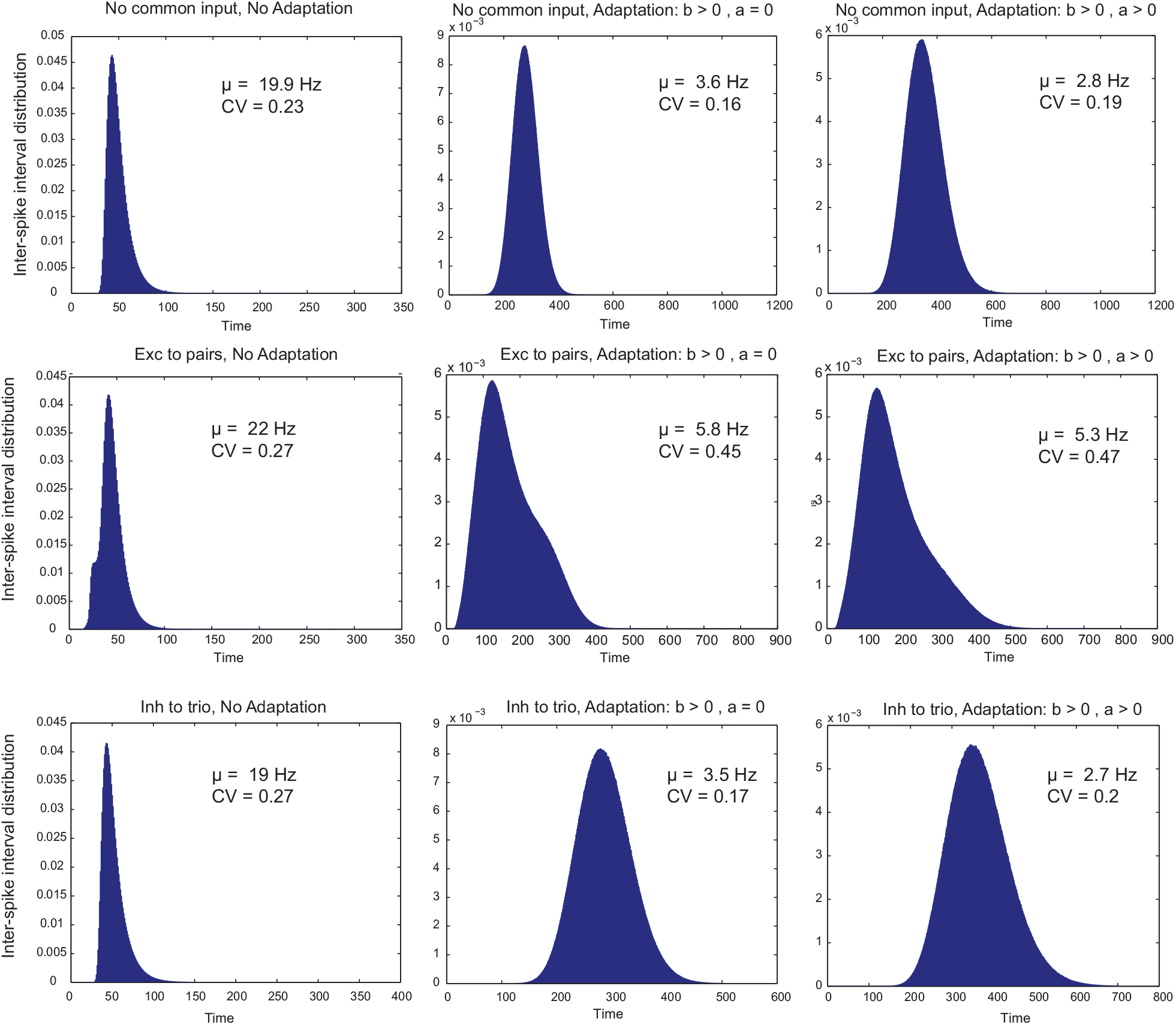
Inter-spike interval (ISI) distributions of postsynaptic neurons for motifs of excitatory-to-pairs and inhibitory-to-trio in the presence and absence of adaptation. The first row shows the ISI when there is no common (signaling) input and the postsynaptic neuron receives mean input which sets the voltage very near to the threshold in addition to a certain level of noise. In the presence of adaptation, when there is no common (signaling) input, the firing rate of postsynaptic neuron decreases, compared to no adaptation case. In the presence of common inputs under excitatory-to-pairs and inhibitory-to-trio motifs, the adaptation decreases the firing rate (second and third row). However, the adaptation increases the coefficient of variation (CV) when excitatory inputs are given to pairs, but decreases the CV in the inhibitory-to-trio case. Fixed parameters are *A/τ*_*m*_*V*_*θ*_ = 0.1, Δ = 10 ms, 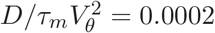, and *λ* = 5 Hz.

**Fig S6.**
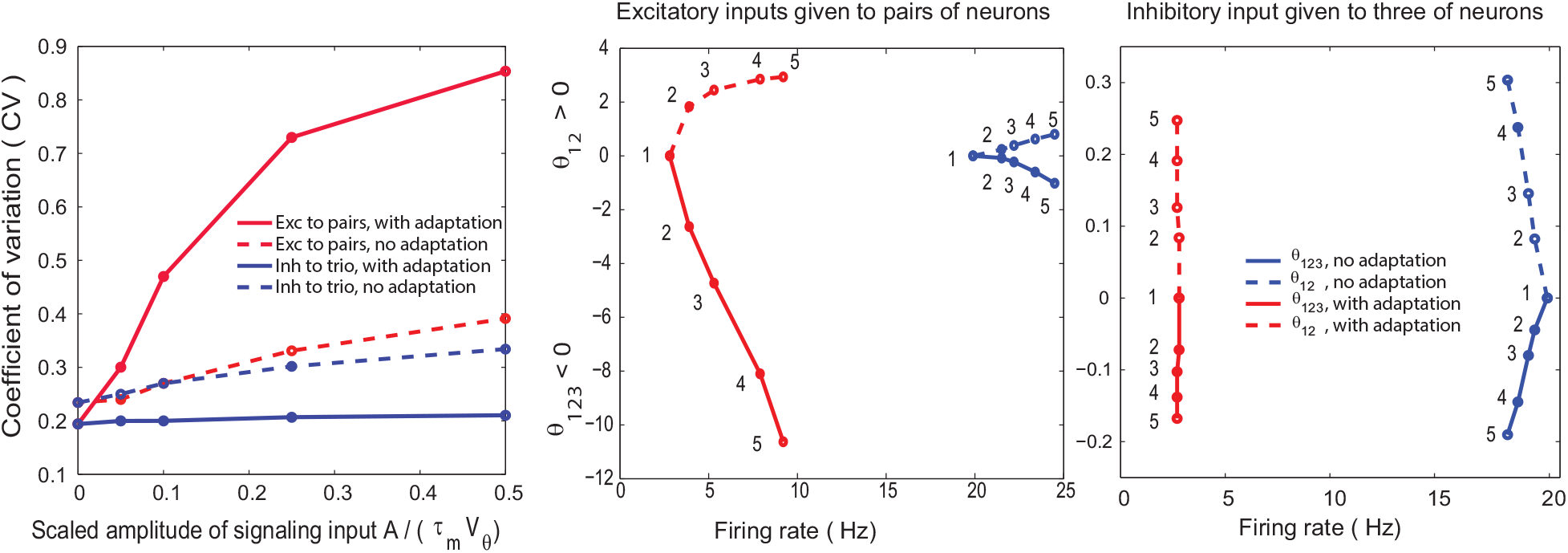
Adaptation can change the coefficient of variation (CV) and interaction parameters in both excitatory-to-pairs and inhibitory-to-trio motifs. (Left) By increasing the amplitude of excitatory inputs given to pairs of neurons, CV can reach high values with the adaptation (red solid line) compared to without the adaptation (left, red dashed line). The adaptation under the inhibitory-to-trio motif decreases the CV (blue lines). (Middle and right) The graph shows *θ*_12_ in the positive domain of the ordinate and *θ*_123_ in the negative domain, as a function of the neurons’ firing rate for excitatory-to-pairs (middle) and inhibitory-to-trio (right) motifs. The red lines present the interactions in the presence of adaptation while the blue is for no adaptation. The numbers 1, 2, 3, 4, 5 assigns to the amplitude of common inputs *A* = 0, 10, 20, 50, 100, respectively. The firing rate is decreased in the presence of adaptation for both excitatory-to-pairs and inhibitory-to-trio motifs (compare each number in blue and red graphs) and even when there is no common input (compare number 1 in blue with red graphs). Nevertheless, *θ*_12_ and *θ*_123_ of excitatory-to-pairs motif are increased for each amplitude of common inputs (middle, compare each number in blue and red graphs). The inhibitory-to-trio motif with adaptation induces smaller interactions, compared with the non-adaptation case (right, compare each number’s interaction in blue with red). Fixed parameters are Δ = 10 ms, 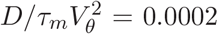, *τ*_*m*_ = 10*ms, V*_*θ*_ = 20 and *λ* = 5 Hz.

### SV: Does mixing motifs, excluding the excitatory-to-pairs motif, can generate strong negative triple-wise interactions?

We have four architectures, common inputs given to pairs or trio of postsynaptic neurons with either common excitatory or common inhibitory inputs. Based on our analysis in this paper about common input and also recurrent activity, we know that the motif of excitatory-to-pairs if exists, can induce strong interactions of *θ*_12_ *>* 0 and *θ*_123_ *<* 0. However, we can ask the following question: By mixing the other motifs (i.e, inhibitory-to-trio, excitatory-to-trio and inhibitory-to-pairs), can we reach the large magnitude of *θ*_12_(*>* 0) and *θ*_123_(*<* 0)? For this purpose, we mix the three motifs, two by two to see whether such a case exists. We mix two motifs with firing rates of *λ*_1_ = 5*Hz* and *λ*_2_ = 5*Hz* for a range of common input’s amplitude in each motif (i.e. from *A* = 0 to *A* = 50) (Fig. S7). The result for mixing excitatory input to trio with inhibitory input to trio is shown in the left panel. Although the inhibitory-to-trio architecture induces negative triple-wise interactions, we observed mostly positive triple-wise interactions when it is mixed with excitatory-to-trio. The only region that the mixing triple-wise interaction are negative is under weak or near zero amplitudes of excitatory inputs given to trio (Fig. S7, left). The mixture of excitatory-to-trio and inhibitory-to-pairs induce positive triple-wise interactions (Fig. S7, middle) as expected from its each individual motif. The result for mixing inhibitory input given to trio and inhibitory input given to pairs is shown in the right panel. Although the inhibitory-to-pairs induces positive triple-wise interactions, when it is mixed with inhibitory-to-trio, for most amplitudes of mixing, negative triple-wise interactions are observed (Fig. S7, right). The interactions induced in this mixing is in the range of inhibitory-to-trio’s interaction which is not strong and does not interfere with excitatory-to-pairs’ range of interactions. The triple-wise and pairwise interactions in each case, are saturated for a specific amplitude of common inputs.

**Fig S7.**
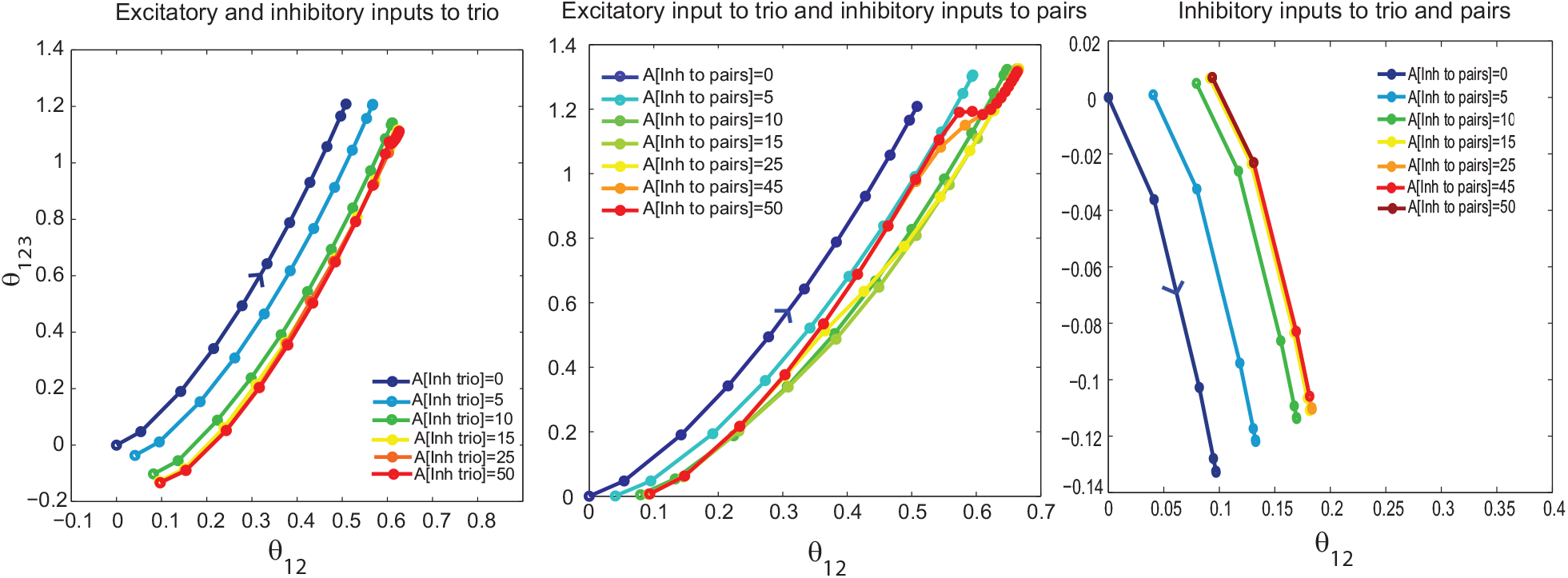
Mixing the motifs of excitatory-to-trio, inhibitory-to-trio, and inhibitory-to-pairs. The result of interactions are shown for the mixture of excitatory and inhibitory inputs each given to trio (left), excitatory-to-trio with inhibitory-to-pairs (middle), and inhibitory-to-trio with inhibitory-to-pairs (right). In the mixture, the amplitude of common input in each motif varied from 0 (i.e. the motif is inactive) to 50 (arrow shows increasing the amplitude in each panel), and the interactions (*θ*_123_ and *θ*_12_) are calculated for different amplitudes of motifs. Along each colored line (following the arrow in each panel), one motif’s amplitude is fixed while the other mixed motif’s amplitude varied from 0 to 50. For example in the left panel, for each colored line, the amplitude of inhibitory-to-trio is fixed (see color box), while the amplitude of excitatory-to-trio has changed along the line from 0 to 50. The color box shows in each panel which motif has the fixed amplitude. Fixed parameters are Δ = 10 ms, 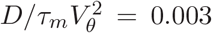, *τ*_*m*_ = 10*ms, V*_*θ*_ = 5mV, *λ*_1_ = 5 Hz and *λ*_2_ = 5 Hz.

### SVI: Why the information-geometric measure is suitable for quantifying pairwise and higher-order interactions

In this study, we analyze the pairwise and higher-order (i.e., triple-wise) statistical dependency among the two and three neurons by using the information-geometric measure that is obtained from the neurons’ interactions expressed in the form of exponential family distribution (Eqs. 13 and 15) (Amari, 2009b; Martignon et al., 2000; Nakahara and Amari, 2002). Here we explain why we employ this measure instead of the other alternatives.

For a system with two neurons (Eq. 13), the information-geometric measure *θ*_12_ of the pairwise interaction expresses the dependency of the two neurons, similarly to the Pearson’s correlation coefficients computed for the binary patterns. Although the zero values of both measures indicate that the two neurons are uncorrelated, their mathematical expressions are different. The information-geometric measure of the pairwise interaction can take an unconstrained real-value whereas the Pearson’s correlation coefficients are bounded in −1 and 1.

It is known that the information-geometric measures have a statistically suitable property to extract the pairwise and higher-order dependency of the binary representation of neural activity (Amari, 2009b; Martignon et al., 2000), compared to conventional measures such as Pearson’s correlations, pairwise and triple-wise cross-correlations (Shlens et al., 2006), cumulants (Staude et al., 2010), and mutual information (Amari, 2009b; Martignon et al., 2000; Ohiorhenuan and Victor, 2011). For example, in the two neurons’ system, the estimation of *θ*_12_ from the data is not affected by estimation of the activity rates of individual neurons. This property does not hold for the other measures such as the classical Pearson’s correlation coefficient (Amari, 2009b). Namely, the simultaneous estimation of the firing rates and the neurons’ dependency measured by the Pearson’s correlation coefficients are correlated. On the contrary, the information-geometric measure of the correlation *θ*_12_ is independent of the estimated firing rates of neurons. In this sense, it quantifies the genuine interaction that we cannot obtain from activities of individual neurons, rather that can be inferred only if we simultaneously observe the two neurons. In information geometry, this property is known as the orthogonality of the interaction to the activity rates of neurons.

For a system with three neurons, the interaction among the three neurons (triple-wise interaction, *θ*_123_) is obtained by Eq. 15. Similar to the information-geometric measure of the pairwise interaction, the triple-wise interactions estimated from the spike data are not correlated with the lower-order statistics, namely activity rates of individual neurons and joint activity rates of the two neurons. It quantifies a genuine triple-wise activity of the three neurons that can be inferred only if we observe the three neurons simultaneously.

The meaning of the non-zero triple-wise interaction becomes clear if we compare it with the neural activity that expresses no triple-wise interaction. Suppose the triple-wise interaction is fixed at zero. In that case, the model corresponds to the maximum entropy model that is derived as the most unstructured model when the individual firing rates and joint firing rates of pairs of neurons (equivalently, pairwise correlations) are given. Such a model produces the joint activities or in-activities of all three neurons (i.e., the pattern ‘111’ or ‘000’) with non-zero probabilities, but they occur as a chance coincidence given the neurons’ firing rates and pairwise correlations. The non-zero information-geometric measure of the triple-wise interaction *θ*_123_ precisely measures the deviation of the joint activity or inactivity of the three neurons from this expected level. If it is positive, the three neurons are jointly active more frequently than the chance level expected from their activity rates and pairwise correlations. If it is negative, the neurons are simultaneously silent more frequently than the chance level.

Comparison of the other methods with the information-geometric measure to calculate the triple-wise (or higher-order) dependency, such as covariance and mutual information (or Kullback-Leibler (KL) divergence), are also summarized in (Amari, 2009b). The former method (covariance) depends on firing rates, and the latter KL-divergence cannot show the sign of correlation. Ohiorhenuan and Victor compared the KL divergence and information-geometric measure, which pointed to the superiority of the information-geometric measure in practice (Ohiorhenuan and Victor, 2011): It can distinguish whether the interactions are less or more synchronous comparing with the chance level while the KL divergence can measure the magnitude only. The information-geometric measure is also easier to calculate the confidence bound, and it was reported that it is robust to spike-sorting errors (Ohiorhenuan and Victor, 2011). In summary, although information-geometric measure needs data to be binarized, its statistical and practical advantages make it suitable for measuring the dependency of neural population.

### SVII: How geometrical constraints modify the probability of finding various microcircuits: A simple Monte Carlo approach

We wrote a simple Matlab code, using 10^8^ times uniform random number generator. It put points on the XY-plane, in a 2*r* × 2*r* square, with *r* = 150*µm*. Then, excluding any point further from the origin (i.e. at *x* = 0 and *y* = 0) than *r*, we simply have a pool of *random points* all sitting in a circle of radius r= 150*µm*. Then, picking each pair (or trio), we verified if two (or three) picked points are closer to each other than the critical length of *l*_c_ = 100*µm*. It simply produces two populations among *all possible choices* of two (or three) points. Out of 4.8 × 10^7^ choices of pairs, we found that the probability of the two points being closer to each other than *l*_c_ = 100*µm* was 0.32; and the probability of finding three points, each closer to two others than 100*µm* was 0.07.

